# Acetyl-CoA carboxylase 1 regulates chondrocyte metabolic plasticity with divergent effects on articular cartilage pathology and growth plate development

**DOI:** 10.64898/2026.11.15.766862

**Authors:** Anupama Binoy, Shadi Moradi, Huanhuan Liu, Grant Davis, Shivakumar R. Veerabhadraiah, Michael J. Jurynec, Martin-Paul Agbaga, P. Christeen Fernando, Mengliang Zhang, Xiaotian Feng, Véronique Lefebvre, Zhaoyang Liu, Vishwajeet Puri, Birgit Schilling, Bent Brachvogel, Shouan Zhu

## Abstract

Osteoarthritis (OA) is a leading cause of disability, yet effective disease-modifying treatments remain unavailable. Metabolic dysfunction is increasingly implicated in OA pathogenesis, although its contributions to cartilage degeneration remain incompletely understood. Here, we investigated the role of acetyl-CoA carboxylase 1 (ACC1, encoded by *Acaca*), a rate-limiting enzyme in *de novo* lipogenesis, in cartilage homeostasis and disease. We found that ACC1 expression was elevated in human and murine articular cartilage during aging and obesity. Cartilage-specific deletion of ACC1 attenuated obesity-associated OA in both male and female mice, reducing cartilage degeneration and pain-related sensitivity without substantially altering systemic metabolic responses to a high-fat diet. ACC1 deletion also reduced pain-related responses following anterior cruciate ligament rupture, with significant structural protection observed predominantly in females. Consistent with these genetic findings, pharmacological ACC inhibition attenuated obesity-exacerbated post-traumatic OA. In contrast, ACC1 deletion during early postnatal development severely disrupted growth plate organization in the long bone, impaired chondrocyte maturation, and reduced longitudinal bone growth, accompanied by subsequent impairment of trabecular bone architecture. Mechanistically, ACC1 deficiency reduced *de novo* lipogenesis and cellular malonyl-CoA levels, accompanied by widespread reductions in protein lysine malonylation. Quantitative malonylome profiling revealed decreased malonylation of key enzymes involved in glycolysis, pentose phosphate pathway, and TCA cycle, including catalytically relevant residues in GAPDH and Aldolase. These changes lead to increased activities of metabolic enzymes, enhanced glycolysis, and extensive remodeling of cellular energy and lipid metabolism. Together, these findings establish distinct roles for ACC1 in articular cartilage homeostasis and growth plate development, highlighting the therapeutic potential of targeting ACC1 in adult OA while underscoring its essential contribution to normal skeletal growth.

## Introduction

Osteoarthritis (OA) is the most common form of arthritis and a leading cause of musculoskeletal disability worldwide, yet no approved disease-modifying therapy is currently available^1,2^. OA is a whole-joint disease involving complex interactions among articular cartilage (AC) , subchondral bone, synovium, and other joint tissues that progressively promote inflammation, cartilage matrix degradation, and loss of joint function^3^. Within AC, chondrocytes are responsible for maintaining extracellular matrix homeostasis through a tightly regulated balance between anabolic and catabolic processes^4^. Increasing evidence indicates that disruption of chondrocyte metabolic homeostasis is an important driver of OA pathogenesis, particularly in the context of major OA risk factors, including aging, obesity, and joint injury.

Altered lipid metabolism has emerged as an important component of this metabolic dysfunction. Obesity and metabolic syndrome are strongly associated with increased risk and severity of knee OA^5–7^, and accumulating evidence suggests that this relationship extends beyond increased mechanical loading. Elevated circulating and tissue lipid levels have been associated with OA development^8–10^, while excess free fatty acids can promote inflammatory responses, matrix degradation, and loss of chondrocyte viability^11^. These observations have generated increasing interest in how fatty acid synthesis, utilization, and remodeling influence cartilage homeostasis. However, the molecular mechanisms linking altered lipid metabolism to chondrocyte dysfunction remain incompletely understood, and whether these metabolic pathways can be therapeutically targeted to modify OA progression remains unclear.

Protein post-translational modifications (PTMs) provide an important mechanism through which the cellular metabolic state can be rapidly coupled to protein function^12–15^. We previously demonstrated that lysine malonylation (MaK), a reversible acyl modification characterized by the addition of a malonyl group to lysine residues, progressively accumulates in AC during aging- and obesity-associated OA^16,17^. Malonyl-CoA, the metabolic donor for MaK^18,19^ is generated during *de novo* lipogenesis (DNL) *via* the action of the enzymes called acetyl-CoA carboxylases (ACCs). ACC1 (ACACA), the predominant cytosolic isoform, produces malonyl-CoA for DNL, whereas ACC2 (ACACB), which is associated with the outer mitochondrial membrane, generates a malonyl-CoA pool that primarily regulates mitochondrial fatty acid oxidation through inhibition of carnitine palmitoyl transferase 1 (CPT1). Thus, malonyl-CoA occupies a unique metabolic position, serving both as an essential intermediate for fatty acid synthesis and as a substrate for protein MaK. Our previous work further demonstrated that ACC1-mediated DNL is active in murine AC^20^ and that ACC1 expression is increased under obesity-associated conditions^21^. These observations led us to hypothesize that ACC1 may function as a metabolic hub linking lipid biosynthesis, protein MaK, and chondrocyte function.

In the present study, we investigated the role of ACC1 in cartilage metabolism and OA pathogenesis using complementary genetic and pharmacological approaches in models of obesity-associated and post-traumatic OA. By integrating transcriptomics, proteomics, malonylation-enriched proteomics, targeted metabolomics, and single-cell RNA sequencing, we examined how ACC1 influences chondrocyte metabolic adaptation and protein regulation. Our findings identify ACC1 as a central regulator of chondrocyte metabolic plasticity through MaK and demonstrate that reducing ACC1 activity protects AC under pathological metabolic and mechanical stress. Unexpectedly, we also uncovered a distinct requirement for ACC1 in cartilage growth plates (GPs), revealing context-dependent functions of ACC1-mediated metabolism in adult AC and endochondral bone development.

## Results

### ACC1 activity drives MaK in chondrocytes

Since ACC-produced malonyl-CoA serves both as an intermediate DNL and as the donor for MaK (Fig. 1a), we first defined which ACC isoform was present in the cartilage tissue. Immunoblot analyses of mouse AC showed a robust amount of ACC1 and minimal or undetectable amount of ACC2, indicating that ACC1 is the predominant ACC isoform present in AC chondrocytes (Fig. 1b). Adding to our previous observation that the ACC1 protein level is increased under obesity-associated conditions^21^, we also found here that the level of ACC1 was markedly higher in aged versus young adult human and mouse AC (Fig. 1c, d).

**Fig. 1:**
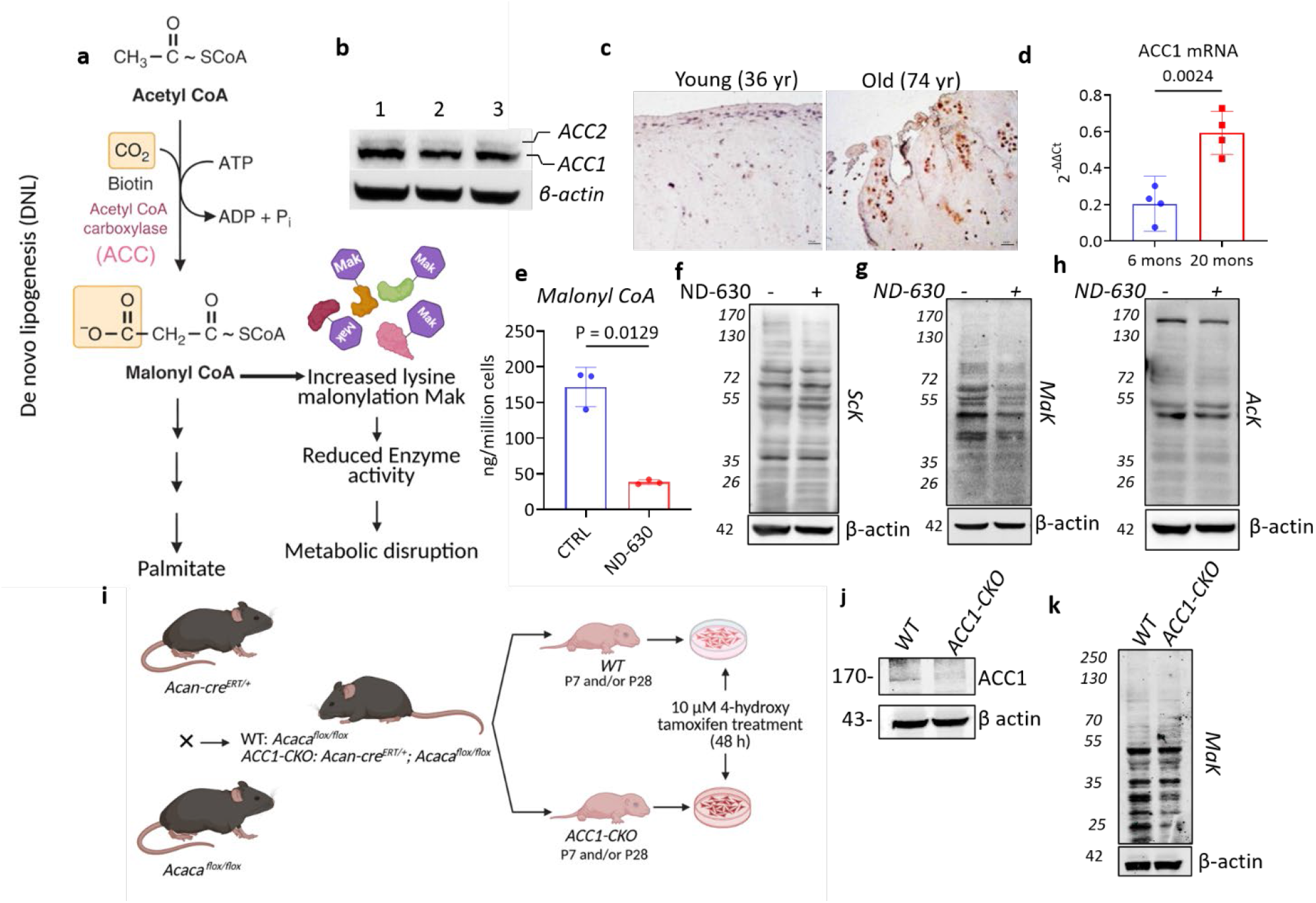
ACC1 is increased in cartilage during aging and obesity conditions, and its inhibition decreases MaK levels. a. Schematic diagram demonstrating the role of ACC in DNL and MaK of proteins. b. Immunoblotting for total ACC in the whole lysate of the AC obtained from WT mice (n=3). c. Immunohistochemical (IHC) staining for ACC1 in human AC tissues from young and old donors. Magnification 10×, (n=3). d. Relative mRNA expression of ACC1 in the AC of 6-month-old and 20-month-old mice (n=4). e. Bar graph showing decreased production of malonyl CoA with ND-630 treatment in primary chondrocytes compared to control (n=3). (f-h). Immunoblotting with Anti-Succinyl lysine Antibody (Sck), g. Anti-Malonyl lysine antibody (Mak), h. Anti-Acetyl lysine antibody (Ack) in primary chondrocytes with or without ND-630 treatment (n=3). i. Schematic diagram showing the crossbreeding strategy to generate *ACC1-CKO* mice. Primary chondrocytes were isolated from the knee joint of Postnatal (P)7 pups (which contain undifferentiated chondrocytes from both AC and GP) and AC of P28 *WT* and *ACC1-CKO* pups (since at the P28 stage there is a clear distinction between AC and GP cartilage^22^). These cells were then used for further experiments. j. Immunoblot showing inhibition of ACC1 in primary chondrocytes isolated from *ACC1-CKO* compared to WT (treated with 4-HT for 48 h to induce cre-mediated knockdown of ACC1), (n=2). k. Immunoblotting with anti-MaK antibody in whole lysates of primary chondrocyte cultures from *WT* and *ACC1-CKO* mice (treated with 10 µM 4-HT), (n=2). Two-group comparisons were performed using an unpaired Student’s T-test with Welch’s correction. p-values < 0.05 were considered statistically significant.

We next tested whether ACC activity regulates MaK in chondrocytes. Primary murine chondrocytes treated with the pan-ACC inhibitor ND-630 (5 nM, 48 h) showed reduced cellular malonyl-CoA (Mean difference of -133.4 ng) level relative to vehicle-treated controls (Fig. 1e). ACC inhibition had little effect on overall lysine succinylation (ScK) (Fig. 1f) but caused a marked reduction in global MaK (Fig. 1g). ND-630 also altered lysine acetylation (AcK) in a selective rather than global manner, with differential changes in proteins of approximately 40 kDa (Fig. 1h). Together, these findings identified ACC activity as a major determinant of MaK in primary chondrocytes, most likely through control of intracellular malonyl-CoA availability.

To define the specific contribution of ACC1, we generated mice with a cartilage-specific *Acaca* knockout by crossing *Acan^CreERT^*^2^*^/+^*mice with *Acaca^flox/flox^* mice, yielding *Acan^CreERT2/+^; Acaca^flox/flox^* (*ACC1-CKO*) animals (Fig. 1i). Primary articular chondrocytes isolated from neonatal knee cartilage were treated *in vitro* with 4-hydroxytamoxifen (4-HT, 10 μM, 48 h) to induce *Acaca* inactivation. Immunoblotting confirmed efficient depletion of ACC1 in *ACC1-CKO* chondrocytes relative to wild-type (WT-*Acaca^flox/flox^*) (Fig. 1j). Consistent with the pharmacological inhibition studies, ACC1 deletion was accompanied by a clear reduction in global MaK (Fig. 1k).

Collectively, these data show that ACC1 expression and enzymatic activity are required to sustain protein MaK in primary chondrocytes, supporting a model in which ACC1-dependent malonyl-CoA production may link metabolic state to post-translational regulation in cartilage *in vivo*.

### Deletion of ACC1 in cartilage protects against obesity-associated knee joint degeneration

To determine whether ACC1 contributes to obesity-associated OA, we administered tamoxifen to WT and *ACC1-CKO* mice at 6 weeks of age *via* intraperitoneal (IP) injections and subsequently fed these mice for 25 weeks with either a low-fat diet (LFD) or a high-fat diet (HFD) (Fig. 2a). Efficient ACC1 deletion in *ACC1-CKO* cartilage was confirmed by immunoblotting and immunohistochemistry, with minimal effect in non-cartilaginous tissues such as liver (Fig. S1a– c). Within each diet group, WT and *ACC1-CKO* mice showed similar body weight gain, fasting glucose, fat mass, and lean mass in both males and females (Fig. S1d–g, l–o). A modest metabolic difference was observed in females, in which *ACC1-CKO* mice displayed improved glucose clearance under HFD conditions, reflected by a lower glucose excursion and reduced area under the curve (Mean difference of 655 units) during glucose tolerance testing; this effect was not evident in males (Fig. S1h–k). Together, these data indicated that cartilage-specific ACC1 deletion does not substantially alter the systemic metabolic consequences of HFD, allowing subsequent joint phenotypes to be interpreted primarily as effects of ACC1 loss in cartilage.

**Figure 2:**
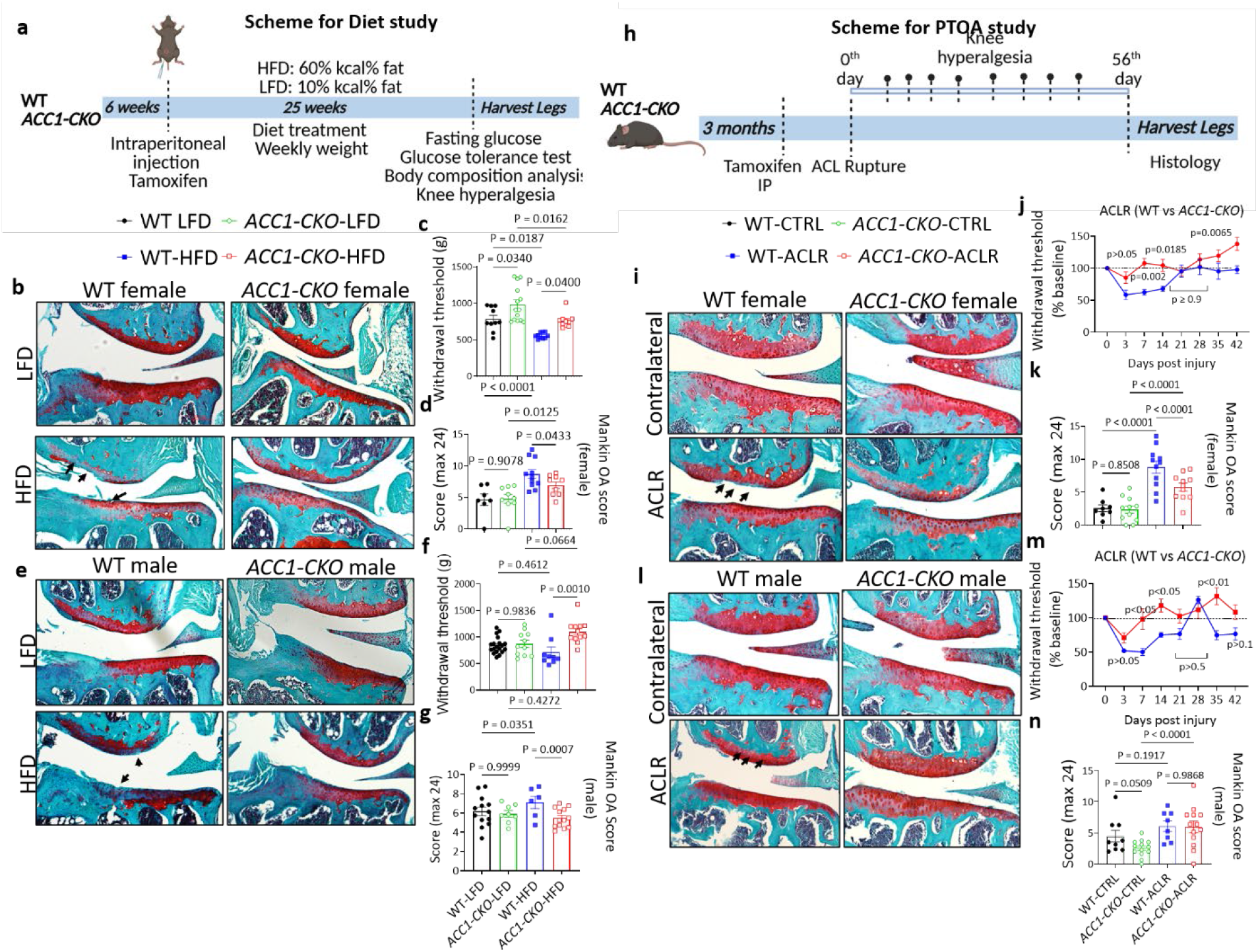
Cartilage-specific disruption of ACC1 protects mice from developing obesity-associated OA and PTOA. a. Schematic description of experimental design to study the effect of ACC1 gene knockdown on obesity associated OA, b, e. Fast green and Safranin O staining of knee joint obtained from female (b), (n=7-10 for WT LFD, n=8-14 for *ACC1-CKO* LFD, n=11 for WT HFD and n=9-10 for *ACC1-CKO* HFD) and male (e), (n=12-19 for WT LFD, n=8-13 for *ACC1-CKO* LFD, n=6-9 for WT HFD, and n=12-13 for *ACC1-CKO* HFD) WT and *ACC1-CKO* mice fed on LFD and HFD. Hyperalgesia measurements, & Mankin OA score for the female (c-d) and male (f-g) mice in the diet study. h. Schematic description of the experimental design to study the effect of ACC1 gene disruption on PTOA. i, l. Fast green and Safranin O staining of knee joints from female (i), (n=10 for WT-contralateral, n=12 for *ACC1-CKO*-contralateral, n=8 for WT-ACLR, and n=12 for *ACC1-CKO*-ACLR) and male (l), (n=9 for WT-contralateral, n=9 for *ACC1-CKO*-contralateral, n=9 for WT-ACLR, and n=10 for *ACC1-CKO*-ACLR) WT and *ACC1-CKO* mice; the upper panel shows the contralateral leg, and the lower panel represents ACLR leg. Hyperalgesia measurements, & Mankin OA score, for the female (j-k) and male (m-n) mice in the PTOA study. Data are expressed as the mean ± SEM. Four-group comparisons were evaluated using two-way ANOVA followed by Tukey’s multiple comparison test. Adjusted p-values < 0.05 were considered significant.

We next assessed whether ACC1 deletion altered lipid accumulation within the joint. Oil Red O staining showed that HFD promoted lipid accumulation in chondrocytes in control mice, and this response was markedly attenuated in *ACC1-CKO* mice (Fig. S1p, q). Furthermore, histological analysis of Safranin O/Fast, Green-stained knee sections showed that HFD-fed WT mice of both sexes developed substantial cartilage damage, proteoglycan loss, and prominent osteophyte formation, particularly at the medial tibial plateau. By contrast, *ACC1-CKO* mice on HFD exhibited less cartilage surface fibrillation, matrix loss, and osteophyte formation (Fig. 2b, e). Under LFD conditions, joint architecture was largely preserved in both genotypes, although *ACC1-CKO* male mice showed milder superficial lesions than WT controls. Together, histological findings indicated overall lower OARSI and Mankin scores for HFD*-*fed *ACC1-CKO* mice than their WT counterparts in both sexes (Fig. S2a, Fig. 2d, and S2b, Fig. 2g,). In females, HFD-fed *ACC1-CKO* mice exhibited a significant mean reduction in OARSI scores of 0.7847 units compared to WT HFD mice (averaging 0.4994 vs 1.284 respectively; p<0.0001), as well as a marked mean decrease in Mankin scores of 1.833 units relative to WT HFD controls (averaging 5.481 vs 7.654 respectively; p=0.0433). In males, HFD-fed *ACC1-CKO* mice similarly showed significantly lower OARSI scores compared to their WT HFD counterparts, reflecting a mean reduction of 0.4477 units (averaging 0.7869 vs 1.235, respectively; p=0.0010), alongside a substantial mean reduction in Mankin scores of 2.174 units (averaging 5.481 vs 7.654, respectively; p=0.0007). Reductions were most apparent in subcomponents reflecting cartilage structural damage, osteophyte formation, and tidemark duplication, whereas other features, including Safranin O staining and hypertrophic chondrocyte scoring, were not significantly affected (Fig. S2a, b). Thus, loss of ACC1 in cartilage mitigates the structural manifestations of obesity-associated knee OA.

Because pain-related behavior is a clinically relevant consequence of OA, we also measured knee hyperalgesia using pressure application measurement testing^23^. *ACC1-CKO* mice exhibited higher withdrawal thresholds (significant mean increase of 203.5g in female and 382.5g in male) than WT mice, particularly under the HFD condition, indicating reduced pain sensitivity despite comparable systemic metabolic stress (Fig. 2c, f). These findings suggest that ACC1 contributes to cartilage structural degeneration and consequently to obesity-associated joint pain.

To investigate possible underlying mechanisms, we modeled obesogenic stress in primary WT chondrocytes by exposure to glucose, insulin, or the combination of glucose plus insulin^24^. Combined glucose and insulin treatment decreased inhibitory ACC phosphorylation, consistent with increased ACC activity, and concomitantly increased MaK. Pharmacological ACC inhibition with ND-630 blunted this increase in MaK, indicating that obesogenic stimulation engages ACC-dependent malonyl-CoA metabolism in chondrocytes (Fig. S1r, s). We next examined the expression of cartilage extracellular matrix anabolic genes. Glucose plus insulin markedly reduced *Col2a1* and *Sox9* expression, and this effect was partially rescued by co-treatment with ND-630, whereas the effects on *Acan* expression were limited (Fig. S1t-v). These data supported the idea that ACC activity contributes at least in part to suppression of cartilage anabolism under metabolically stressful conditions.

Collectively, these data identified cartilage ACC1 as a local driver of HFD-induced lipid accumulation, joint degeneration, and pain-related behavior, and support ACC1-dependent lipid metabolism as a mechanistic link between obesity and OA.

### Cartilage-specific ACC1 deletion attenuates post-traumatic OA in a sex-dependent manner

We next asked whether cartilage-specific loss of ACC1 also protects against post-traumatic OA (PTOA). To minimize potential confounding effects of ACC1 deletion on skeletal development, recombination was induced in adult mice at 3 months of age. PTOA was then induced by non-invasive rupture of the anterior cruciate ligament (ACL) in the left knee^25^, while the contralateral limb served as an internal control (Fig. 2h).

Knee hyperalgesia was assessed before injury and longitudinally thereafter (days 3, 7, 14, 21, and weekly through day 56), with values normalized to pre-injury baseline. Following ACL rupture (ACLR), both WT and *ACC1-CKO* mice of both sexes exhibited a marked reduction in withdrawal threshold at the early post-injury time point, dropping to 58.43% (female), 51.90% (male) for WT ACLR vs 85.24% (female), 70.93% (male) for *ACC1-CKO* ACLR, indicating acute pain sensitization at day 3. However, after this initial decline, *ACC1-CKO* consistently showed higher withdrawal thresholds than WT mice (Fig. 2j, m). Specifically, female *ACC1-CKO* mice demonstrated significant attenuation compared to WT females, with significant group differences observed at key time points such as day 7 (mean difference of -45.37%), day 14 (mean difference of -36.52%) and day 42 (mean difference of -40.32%). Similarly, male *ACC1-CKO* exhibited significantly higher normalized withdrawal thresholds relative to injured WT males at day 7 (mean difference of -47.77%), day 14 (mean difference of -42.72%) and day 35 (mean difference of - 56.89%), suggesting that loss of cartilage ACC1 attenuates persistent injury-associated pain across sexes.

We next examined structural joint degeneration at the study endpoint by histological analysis. ACLR induced substantial cartilage degeneration in both genotypes, characterized by surface fibrillation, deep cleft formation, and loss of proteoglycan staining. However, the severity of these changes was lower in *ACC1-CKO* mice relative to WT controls, with the protective effect being more evident in females than in males (Fig. 2i, l). Summing up these observations, ACL injury increased both OARSI and Mankin scores across genotypes and sexes, but overall histological protection was largely restricted to female *ACC1-CKO* mice, which showed significantly lower total OARSI scores (mean difference of 0.32 units), and Mankin scores (mean difference of 3.023 units) than injured WT females (Fig. S2c and Fig. 2k). By contrast, injured male *ACC1-CKO* mice did not exhibit significant reductions in overall OARSI or Mankin scores relative to injured WT males (Fig. S2d and Fig. 2n).

Analysis of individual Mankin subcomponents further indicated that not all pathological features were equally affected by ACC1 loss. Osteophyte formation and hypertrophic chondrocyte scores were not significantly altered, even in female mice, suggesting that ACC1 deletion preferentially mitigates selected aspects of post-injury cartilage degeneration rather than uniformly suppressing all features of PTOA pathology (Fig. S2c, d).

Together, these findings indicate that cartilage-specific ACC1 deletion reduces pain-related responses after joint injury and confers partial structural protection against PTOA, with the histological benefit being more pronounced in female mice. These data further support ACC1-dependent metabolic regulation as a contributor to OA pathogenesis across both obesity-associated and injury-induced settings, while also revealing a sex-dependent component in the response to ACC1 loss.

### Loss of ACC1 reduces malonylation of central carbon metabolic enzymes and rewires chondrocyte metabolism

Because ACC1 generates malonyl-CoA, the metabolic donor for protein MaK^26,27^, and we have observed ACC1 inhibition decreased MaK, we next asked how ACC1 deletion alters the chondrocyte malonylome. Quantitative MaK profiling of WT and *ACC1-CKO* primary chondrocytes identified 754 malonylated peptides corresponding to 513 malonylation sites (Fig. 3a, b, Supplementary excel data 1). Although many modified peptides showed comparable abundance between genotypes, a substantial subset exhibited reduced MaK in *ACC1-CKO* chondrocytes with many proteins falling below the 1X threshold, consistent with decreased availability of ACC1-derived malonyl-CoA (Fig. 3b). Pathway enrichment analysis of hypomalonylated proteins revealed a striking concentration within central carbon metabolism, including glycolysis/gluconeogenesis, pyruvate metabolism, the pentose phosphate pathway (PPP), the tricarboxylic acid (TCA) cycle, and amino acid biosynthesis (Fig. 3c).

**Fig. 3.**
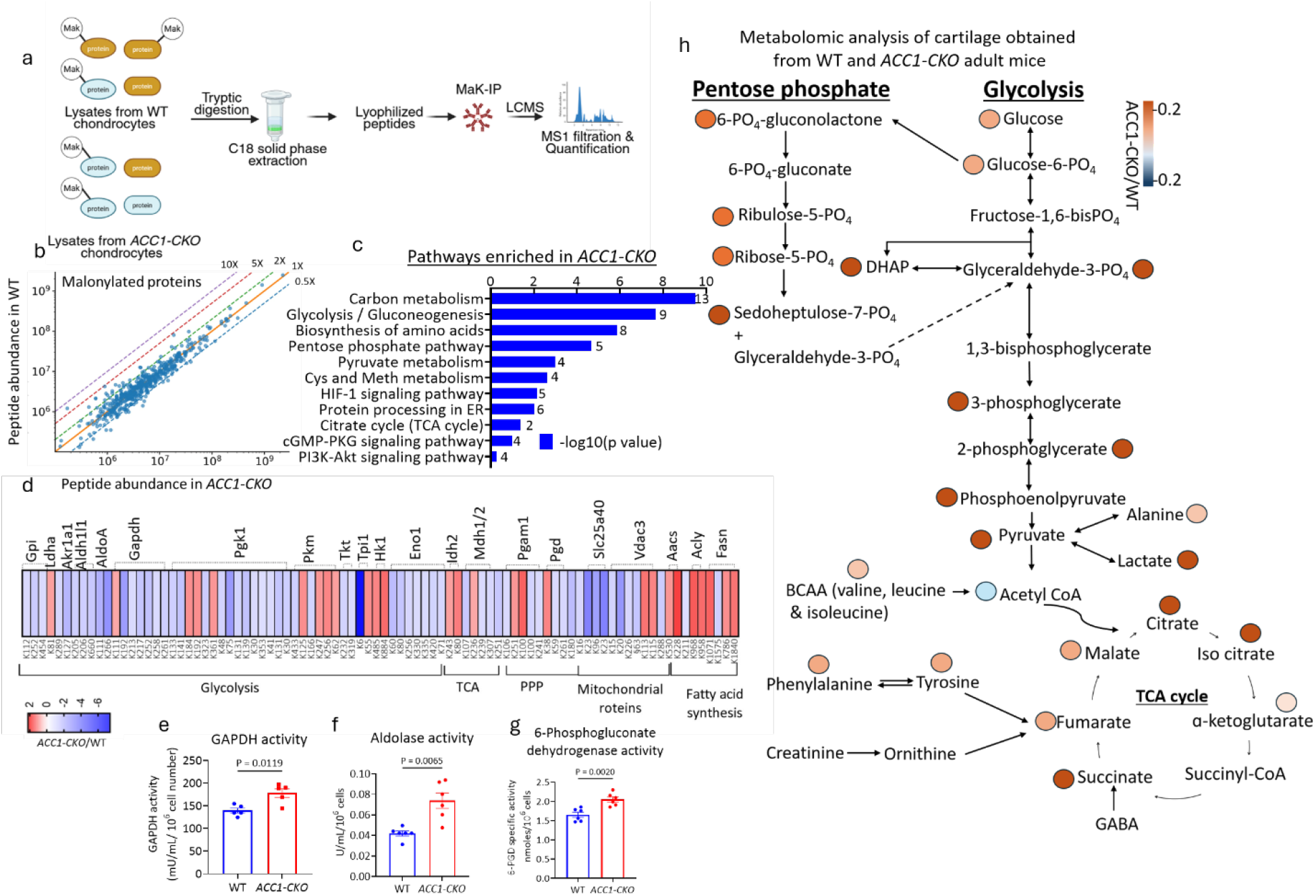
Metabolic rewiring in the AC of *ACC1-CKO* is influenced by the level of malonylation of key enzymes in the central carbon metabolism pathway. a. The workflow illustrates the proteomic strategy used to identify and quantify MaK proteins in chondrocytes; (n=4) b. The scatter plot compares the relative abundance of malonylated peptides between WT and *ACC1-CKO* samples, quantified by LC–MS/MS. c. Pathway enrichment analysis on differentially less malonylated proteins in *ACC1-CKO* cartilage in comparison to WT. Numbers at the end of each bar denote the number of malonylated proteins mapped to each pathway. d. A heat map showing the site-specific malonylation pattern for selected enzymes from pathways such as glycolysis, the TCA cycle, and the PPP, as well as mitochondrial proteins and proteins associated with fatty acid synthesis, e. Enzymatic activity of GAPDH enzyme; (n= 6) f. Enzyme activity of Aldolase A; enzyme (n= 6), g. Specific enzyme activity of 6-phosphogluconate dehydrogenase; (n=6) (enzyme activities were normalized to 10^6^ primary chondrocytes in all assays). h. central carbon metabolic pathway consisting of glycolysis, the TCA cycle, and the PPP, showing normalized fold changes in metabolite peak areas from the targeted metabolomics analysis (n=4 per group). The gradient from red to blue in panel d and h indicates the ratio of integrated peak area of each protein/metabolite abundance in *ACC1-CKO* (numerator) vs WT (denominator) groups. Data are expressed as the mean ± SEM. Two-group comparisons were evaluated using unpaired Student’s t-test with Welch’s correction.

Mapping individual MaK sites onto representative metabolic proteins further revealed extensive but non-uniform remodeling across glycolytic, PPP, and TCA-cycle enzymes, as well as mitochondrial and fatty acid synthesis proteins (Fig. 3d). Importantly, different sites, including multiple sites within the same protein, did not change uniformly, indicating selective remodeling of the malonylome rather than a simple global reduction in MaK. Several key glycolytic enzymes, including GAPDH and PGK1, contained sites with markedly reduced malonylation in *ACC1-CKO* chondrocytes, suggesting that ACC1-derived malonyl-CoA may directly couple lipid metabolism to regulation of central carbon metabolic proteins. We next asked whether this remodeling of MaK was associated with altered activity of the affected metabolic pathways. *ACC1-CKO* chondrocytes exhibited increased activities of GAPDH (mean difference of 38.40mU/mL) and aldolase A (mean difference of 0.31U/mL), together with increased activity of the PPP enzyme 6-phosphogluconate dehydrogenase (6PGD; mean difference of 0.3911 nmoles of NADPH; Fig. 3e–g). These functional changes are consistent with an inverse relationship between MaK and the activity of selected central carbon metabolic enzymes, although they do not by themselves establish a direct site-specific effect of malonylation. Consistent with these enzymatic changes, targeted metabolite mapping revealed broad redistribution of intermediates across glycolysis, the PPP, and the TCA cycle, with many metabolites increased in *ACC1-CKO* cartilage (Fig. 3h, supplementary excel data 2). Among glycolytic intermediates, like glucose-6-phosphate/fructose-6-phosphate (Log_2_FC: 1.13; p<0.0001), glyceraldehyde-3-phosphate/ dihydroxyacetone phosphate (Log_2_FC: 2.986; p<0.0001), 3-phosphoglycerate (Log_2_FC: 1.17; p<0.0001), phosphoenolpyruvate (Log_2_FC: 3.336; p<0.0001), and pyruvate (Log_2_FC: 1.58; p<0.0001) were significantly increased. Despite these increases, acetyl-CoA was reduced (Log_2_FC: -0.63; p=0.0012), whereas multiple TCA-cycle intermediates, including citrate (Log_2_FC: 0.955; p=0.0002), isocitrate (Log_2_FC: 0.955; p=0.0002), α-ketoglutarate (Log_2_FC: 0.05; p<0.0001), succinate (Log_2_FC: 1.14; p<0.0001), fumarate (Log_2_FC: 0.36; p<0.005), and malate (Log_2_FC: 0.68; p<0.0001), were maintained or increased. Similarly, PPP intermediates such as 6-phosphogluconolactone (Log_2_FC: 2.412; p<0.0001), ribulose-5-phosphate (Log_2_FC: 2.683; p=0.004), ribose-5-phosphate (Log2FC: 1.044; p<0.0001), and sedoheptulose-7-phosphate (Log_2_FC: 4.686; p=0.009) were also increased in *ACC1-CKO* vs WT group. Several amino acids and related metabolites, including alanine (Log_2_FC: 0.46; p=0.006), phenylalanine (Log_2_FC: 0.12; p<0.0001), and tyrosine (Log_2_FC: 0.27; p<0.0001), branched chain amino acids were moderately increased. Together, this metabolite profile supports extensive redistribution of carbon metabolism following ACC1 loss, characterized by accumulation of glycolytic and PPP intermediates and maintenance of TCA-cycle metabolites despite reduced acetyl-CoA availability^28^. To functionally determine whether the metabolic remodeling associated with ACC1 loss alters cellular energy metabolism, we performed Seahorse analysis in primary chondrocytes. *ACC1-CKO* cells exhibited increased extracellular acidification, with higher basal (mean difference of 5.639 pmol/min) and compensatory glycolysis (mean difference of 10.77 pmol/min), glycoPER (mean difference of 6.552 pmol/min), and post-2-DG acidification (mean difference of 2.006 pmol/min) compared with WT cells (Fig. S3a–e). In contrast, basal mitochondrial respiration (mean difference of-1.159 pmol/min) was largely maintained, whereas maximal respiration (mean difference of -3.898 pmol/min) and spare respiratory capacity (mean difference of -1.717 pmol/min) were reduced in *ACC1-CKO* chondrocytes, with relatively modest effects on ATP-linked respiration (mean difference of - 0.6410 pmol/min) (Fig. S3f–j). Consistent with these changes, the Seahorse energy phenotype map shifted *ACC1-CKO* cells toward a more glycolytic basal state (Fig. S3k). Glucose uptake was also increased in *ACC1-CKO* chondrocytes (mean increase of 50.44 units), further supporting greater reliance on glucose utilization (Fig. S3l). Together with the increased activities of glycolytic and PPP enzymes and the accumulation of central carbon metabolites described above, these findings indicate that ACC1 loss promotes a shift toward glycolytic metabolism while preserving basal mitochondrial respiration, but with reduced capacity to increase oxidative respiration under metabolic stress.

We next examined whether this metabolic adaptation was associated with altered oxidative stress. Basal intracellular ROS levels were similar between WT and *ACC1-CKO* chondrocytes, as assessed by DCFDA fluorescence (Fig. S3m). However, NOX4 expression was reduced in *ACC1-CKO* AC (Fig. S3n), suggesting that the metabolic shift induced by ACC1 loss does not increase basal oxidative burden and may reduce a major source of ROS production in cartilage.

Given chondroprotective effect of ACC1 knockout, we predict that sustained ACC enzymatic activity produces a reciprocal cartilage phenotype. Thus, we examined ACC double knock-in (*ACC-DKI*) mice ^13,29^ with alanine knock-in mutations at ACC1 Ser79 and ACC2 Ser 212 (ACC-DKI) preventing AMPK driven inhibitory phosphorylation, resulting in constitutive ACC activation and malonyl-CoA production that is no longer responsive to AMPK regulation. AC from ACC-DKI mice exhibited greater structural degeneration with increased Mankin (Median difference of -6.500) and insignificantly changed OARSI (Median difference of -1.500) scores compared with WT controls (Fig. S3o, p). Metabolomic profiling of the AC further revealed altered central carbon metabolism, including reduced abundance of several amino acids, particularly branched-chain amino acids, glutamate, histidine, and phenylalanine, whereas changes among glycolytic metabolites were more limited, with glucose-1-phosphate (0.6541 units) and glucose-6-phosphate (0.6652 units) among the metabolites reduced in *ACC-DKI* cartilage (Fig. S3q, r, supplementary excel data 3).

### ACC1 loss induces a metabolically adaptive program in AC chondrocytes

To further investigate the underlying molecular mechanisms, we performed multi-omics experiments. Bulk RNA-sequencing of adult WT and *ACC1-CKO* AC identified 941 upregulated and 608 downregulated genes following ACC1 deletion (Fig. S4a, supplementary excel data 4). Consistent with loss of the rate-limiting enzyme for DNL, genes involved in fatty acid synthesis, elongation, and desaturation, including *Acly, Elovl6, Fads2,* and *Fads3*, had their expression reduced. ACC1 deletion also suppressed inflammatory lipid signaling, with decreased expression of genes involved in arachidonic acid mobilization and eicosanoid synthesis, including *Alox12, Ptgs2,* and several phospholipase A2 family members. In contrast, genes involved in alternative lipid handling and sequestration, including *Fabp3, Acacb, Cidea,* and *Cidec*, had their expression increased (Fig S4b). These changes suggest that disruption of DNL is accompanied by compensatory redistribution and utilization of lipid pools while reducing the activity of pathways associated with pro-inflammatory lipid production. Inflammatory and matrix-associated programs were also selectively remodeled. Expression of *Il17f, Il17b,* and several immune-associated chemokines were reduced, whereas stress-response genes including *Hspa1a* and *Hspa1b* were upregulated (Fig. S4c). Extracellular matrix genes showed a similar selective rather than uniform response: *Acan*, type IX collagen genes like *Col9a1*, *Col9a3*, and *Col10a1* were downregulated, whereas *Col2a1* expression remained relatively unchanged (Fig. S4d). Thus, ACC1 loss does not suppress all features of the articular chondrocyte phenotype but instead shifts matrix and inflammatory programs toward less hypertrophic and less inflammatory states.

A prominent feature of the transcriptomic response was the activation of alternative energy-producing pathways. Multiple glycolytic genes, including *Gapdh, Hk2, Pfkm, Aldoa,* and *Pgam2*, were upregulated together with genes for components of AMPK signaling and genes involved in mitochondrial oxidative metabolism (Fig. S4e, f). GSEA reinforced this pattern, showing positive enrichment of glycolysis/gluconeogenesis, pyruvate metabolism, the TCA cycle, oxidative phosphorylation, AMPK and PPAR signaling, and adipocytokine signaling. Conversely, cytokine– cytokine receptor interaction, IL-17 signaling, ECM–receptor interaction, and DNA replication were impoverished (Fig. S4g). Collectively, the transcriptomic data indicated that loss of ACC1 shifts AC away from lipid synthesis, inflammatory signaling, and proliferation and engages into alternative pathways for energy production and metabolic adaptation.

We next asked whether these transcriptional changes were reflected at the protein level by performing quantitative proteomics in primary WT and *ACC1-CKO* AC chondrocytes (P28). Among 313 significantly altered proteins, 172 were increased, and 141 were decreased following ACC1 depletion (Fig. S5a, supplementary excel data 5). Proteins involved in autophagy and mitophagy, including ULK1, ATG9A, MAP1LC3B, GABARAPL family members, SQSTM1, BNIP3, and LAMP2, robustly increased (Fig. S5b), indicating enhanced engagement of intracellular recycling and quality-control pathways. ACC1-deficient chondrocytes also showed increased amounts of proteins involved in lipid uptake, trafficking, activation, and oxidation, including FABP family members, ACSL3/4, CPT1A, ACOX1/3, and other lipid-remodeling proteins (Fig. S5c). These findings further support a compensatory shift from endogenous fatty acid synthesis toward acquisition, redistribution, and utilization of existing lipid pools.

At the same time, the amounts of proteins involved in cell-cycle progression and DNA replication, including PCNA, MCM2–7, CDK1, CDK2, CCNB1, and CDC20, were broadly reduced, and the same was true for several integrins and pericellular matrix proteins (Fig. S5d, f). Proteins involved in iron handling, glutathione metabolism, and protection against lipid oxidative stress were over-represented (Fig. S5e). Proteomic GSEA consequently revealed positive enrichment of autophagy, mitophagy, PPAR signaling, ether lipid metabolism, cholesterol metabolism, and fatty acid degradation, together with negative enrichment of DNA replication, cell cycle, ribosome biogenesis, mismatch repair, and ECM–receptor interaction (Fig. S5g). Thus, the proteomic data closely matched the transcriptomic findings, defining a state characterized by enhanced metabolic flexibility and cellular quality control coupled to reduced proliferative and anabolic demand.

To integrate these responses, we combined the transcriptomic and proteomic enrichment signatures using network analysis. Three major functional modules emerged (Fig. 4a). Module 1 centered on lipid metabolism and nutrient sensing, linking fatty acid metabolism, PPAR and adipocytokine signaling, AMPK–mTOR pathways, autophagy/mitophagy, and ferroptosis-related processes. Module 2 comprised central carbon metabolic pathways, including glycolysis, the PPP, pyruvate metabolism, and the TCA cycle, and was closely connected to Module 1. In contrast, Module 3 contained DNA replication, cell-cycle, and DNA-repair pathways and was relatively separated from the metabolic modules. This organization suggests a coordinated redistribution of cellular resources in *ACC1-CKO* articular chondrocytes, toward energy production, lipid adaptation, redox control, and cellular maintenance.

**Fig. 4:**
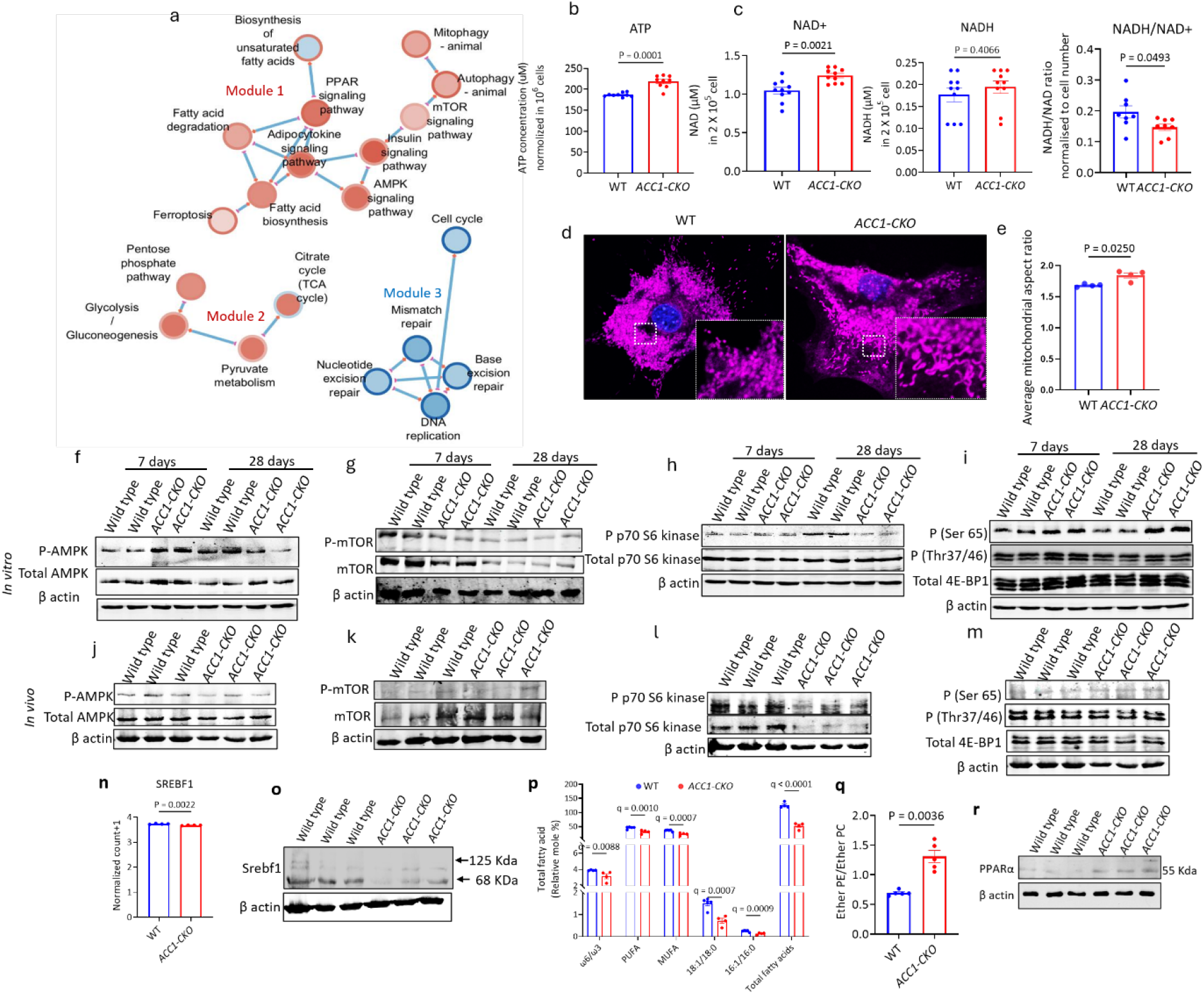
Integrated transcriptomic–proteomic analysis reveals modulation in lipid/nutrient sensing and stress response, central carbon metabolism, and cell fate in ACC1-deficient cartilage. a. Cytoscape-based RNA-protein interaction network showing pathway nodes representing significantly enriched gene sets from transcriptomic (Node fill color depicts RNA NES; red = positive enrichment/upregulated program, blue = negative enrichment/downregulated program; and proteomic (Node border depicting Protein NES; red = positive, blue = negative) GSEA (KEGG mmu pathways). Edges connect pathway pairs with shared leading-edge (“core”) genes, indicating functional coupling, b. Measurement of ATP production in whole lysate of WT and *ACC1-CKO* P7 primary chondrocytes normalized to 10^6^ cells per sample, c. quantification graphs for NAD+, NADH, and NADH/NAD+ ratio in homogenized P7 primary chondrocytes from WT and *ACC1-CKO*, d. Confocal microscopic images of WT and *ACC1-CKO* P7 primary chondrocytes after staining the mitochondria with MitoTracker Deep Red; the inset boxes show the magnified images of the dashed-square area, e. quantification of the tubular and fragmented mitochondria (n=3 per group). Whole lysates of P7 and P28 primary chondrocytes (f, g, h, i) and adult AC tissue (femur condyl and tibial plateau) (j, k, l, m) from WT and *ACC1-CKO* mice were immunoblotted with anti-p-AMPK, total AMPK, p-mTOR, total mTOR, p-p70 S6kinase, total p70 S6kinase, p-(Ser65) and p-(Thr37/46) 4E-BP1 and total 4E-BP1. *Phospho-proteins and their total proteins were probed on separate membranes and were normalized to the respective β-actin as a loading control. n. Normalized count bar plot showing the expression of the *Srebf1* gene in WT and *ACC1-CKO* cartilage tissue, n=4. o. Western blot showing the reduced conversion of premature SREBF1 to mature SREBF1 in *ACC1-CKO* cartilage tissue from adult mice. p. Bar graph depicting the relative mole percent of ω6/ω3 fatty acids, polyunsaturated fatty acids (PUFAs), monounsaturated fatty acids, ratio of 18:1/18:0 fatty acids, ratio of 16:1/16:00 fatty acids, and total fatty acids obtained from WT and *ACC1-CKO* P7 primary chondrocytes (n=4), q. Ratio of relative % of ether PE and ether PC in WT and *ACC1-CKO* P7 primary chondrocytes (n=4), r. Western blot showing the expression of PPAR-α in WT and *ACC1-CKO* AC tissue. Data are expressed as the mean ± SEM. Two-group comparisons were evaluated using unpaired Student’s t-test with Welch’s correction. p-values <0.05 were considered significant. For panel p individual row analysis comparing WT and *ACC1-CKO* were performed using multiple unpaired t-tests without assuming equal SDs, and significant hits were determined using the two-stage step-up FDR method (Q=1%). Statistically significant value was set to p or q<0. 05. see supplementary material Fig 1 uncropped bots.

We next performed functional analyses to determine whether this omics-defined adaptive state was accompanied by improved bioenergetic homeostasis. *ACC1-CKO* articular chondrocytes contained more ATP than WT cells (Mean difference of 31.89 µM) and exhibited increased NAD+ abundance (0.1956 µM) together with a lower NADH/NAD+ ratio (Mean difference of -0.04947 Units) (Fig. 4b, c), consistent with improved bioenergetic homeostasis. MitoTracker staining further showed a more elongated and interconnected mitochondrial network in *ACC1-CKO* chondrocytes, indicative of reduced fragmentation and a shift toward mitochondrial fusion (Fig. 4d, e). Immunoblot analysis showed reduced phosphorylation of AMPK, consistent with diminished energetic stress (Fig 4f, j and Fig. S6a, f). Notably, phospho-mTOR levels also decreased (-1.613 fold) in the *ACC1-CKO* cartilage, which points towards non-independence on AMPK driven regulation^30^ (Fig. 4g, k and Fig. S6b, g). Consistent with mTOR inactivation, decreased phosphorylation of p70 S6 kinase (-0.3994-fold) (Fig. 4h, l and Fig. S6c, h) indicated that ACC1-deficient chondrocytes had restricted anabolic translation, probably to support maintenance of cartilage. Despite this overall suppression of anabolic mTOR signaling, phosphorylation of 4E-BP1 at Ser65 (0.2565-fold) and Thr37/46 (0.2633-fold) increased in *ACC1-CKO* chondrocytes (Fig 4i, m and Fig. S6d, i, e, j), suggesting differential, site-specific regulation of translation. Collectively, these data support a model in which ACC1 loss drives a bioenergetically favorable, stress-adaptive state in chondrocytes characterized by lipid remodeling, autophagy-associated quality control, and reduced anabolic signaling.

### ACC1 deletion suppresses DNL and remodels the chondrocyte lipidome

Consistent with decreased mTOR activity, we observed decreased expression of the *Srebf1* gene in *ACC1-CKO* vs WT cartilage transcriptomics analysis (Fig. 4n), indicating reduced lipogenesis upon ACC1 deficiency. Here, we found that ACC1 deletion reduced formation of the mature SREBF1 transcription factor (Mean difference of 1.875) (Fig. 4o and S6k), which reduced the relative abundance of total fatty acids (Mean difference of 72.65 %) (Fig. 4p) and was thus consistent with overall suppression of DNL. Oil Red O staining of cultured WT and *ACC1-CKO* primary chondrocytes revealed a clear reduction in neutral lipid accumulation following ACC1 deletion (Fig. S6l). This decrease was confirmed by showing that the proportion of lipid-containing particles (LPC), relative to total chondrocyte number, was significantly lower in *ACC1-CKO* cultures (Mean difference of -0.2376) than in WT (Fig. S6m).

In parallel, the omega-6 to omega-3 fatty acid ratio was reduced in *ACC1-CKO* chondrocytes (Mean difference of -0.75%) (Fig. 4p), consistent with a shift toward a lipid composition generally considered less pro-inflammatory. Lipidomic analyses of phosphatidylcholine (PC) and phosphatidylethanolamine (PE) species revealed broad changes in phospholipid composition in *ACC1-CKO* chondrocytes relative to WT cells (supplementary excel data 6). In particular, the ratio of ether-linked PE to ether-linked PC was significantly increased in *ACC1-CKO* chondrocytes (Mean difference of 0.6144) (Fig. S6n, o and Fig. 4q), further supporting a marked reorganization of membrane-associated lipid species after loss of ACC1. This observation was further supported by the overexpression of PPAR-α at the RNA (Mean difference 0.9987) and protein levels in *ACC1-CKO* cartilage tissue (Mean difference of 2.276) (Fig. 4r and Fig. S6p, q, r). Together, these data showed that ACC1 deletion not only reduces overall lipid accumulation and fatty acid synthesis in chondrocytes but also drives selective remodeling of fatty acid and phospholipid pools, consistent with a substantial rewiring of chondrocyte lipid metabolism.

### ACC1 is required for GP integrity and longitudinal bone growth

During histopathological analysis of knee joints from obesity and PTOA studies, we unexpectedly observed that deletion of ACC1 in adult mice promoted epiphyseal GP closure, a non-existent phenomenon in long bones of adult WT mice (Fig. 5a). Consistent with the fact that adult mouse GPs are barely active, this GP closure in mutant mice did not result in any visually obvious difference in overall body size between wild-type (WT) and *ACC1-CKO* animals. This finding suggested that ACC1 may be required for GP integrity, and consequently for longitudinal bone growth in neonatal and juvenile mice^31^. To test this directly, we established a separate experimental paradigm in which tamoxifen was administered by oral gavage to lactating WT dams mated with *ACC1-CKO* males while the pups were at the postnatal days 2 (P2) to P4 (Fig. 5b). Offspring were then analyzed at P14, P28 and P120. *ACC1-CKO* mice of both sexes exhibited reduced naso-tail tip length as early as P14, and this difference persisted with age (Fig. 5c and Fig. S7a). Furthermore, tibial and femoral lengths were reduced in *ACC1-CKO* mice relative to WT controls (Fig. 5d, e and Fig.S7c, d). By contrast, body weight divergence became significant only after P28 (Mean difference of 3.7g) (Fig. S7b), with a reduced fat mass % at both P28 (Mean difference of 3.03%) and P120 (Mean difference of 5.99%) and reduced lean mass % at only P120 (Mean difference of -4.28%) in the *ACC1-CKO* mice (Fig. S7e, f). Safranin O/Fast Green staining of proximal GP at P28 showed that *ACC1-CKO* GPs displayed disrupted columnar organization within the proliferative zone (PZ), with shorter and less orderly chondrocyte columns than in WT mice, but no major change in overall PZ width (Fig. 5f). These abnormal clusters contained more cells than WT columns (mean difference of -86.46 no. of cells), whereas the hypertrophic zone (HZ) was reduced in width (mean difference of 30.02 mm) (Fig. 5g, h).

**Figure 5:**
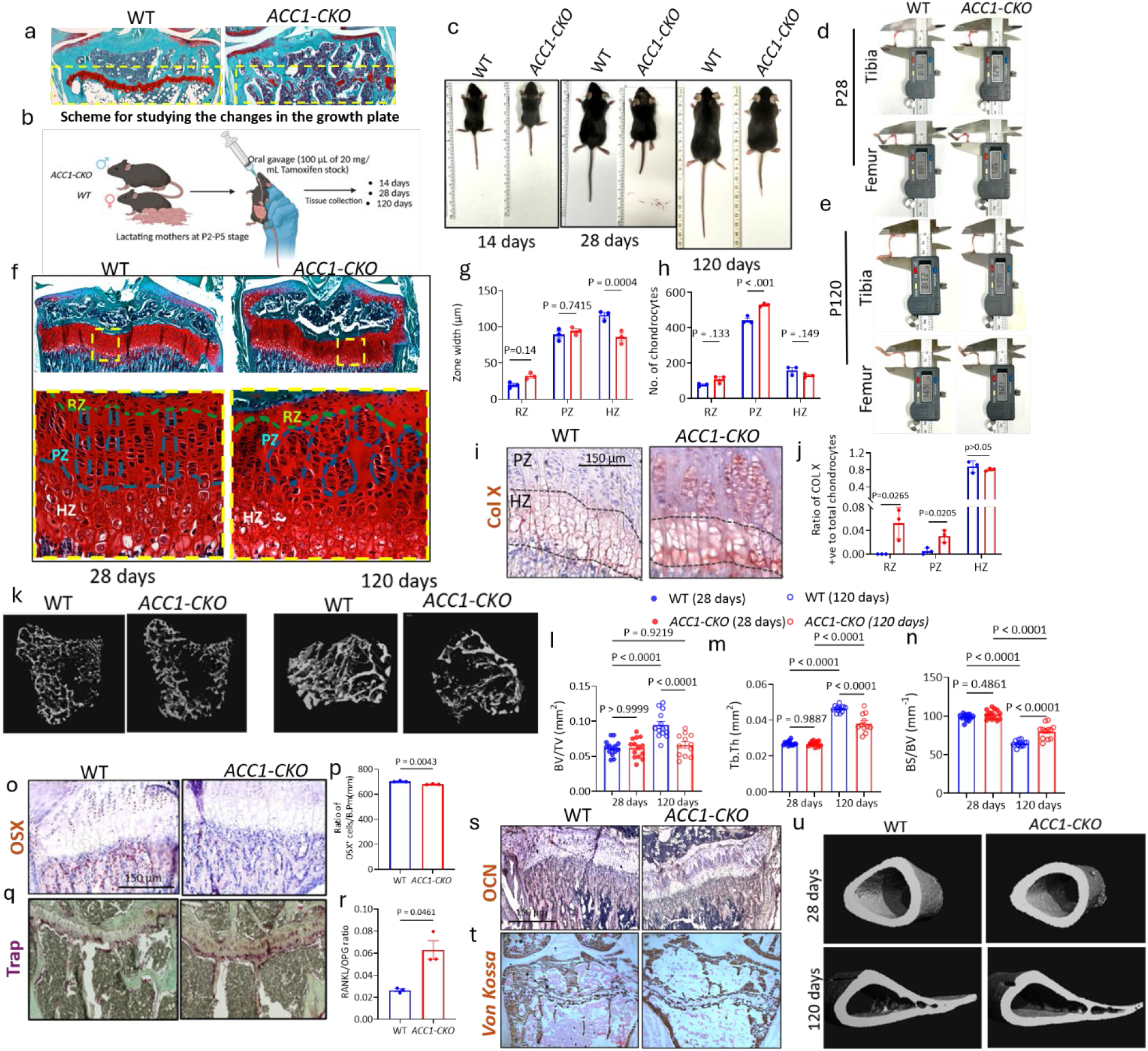
Disruption of ACC1 at neonatal stages affects the growth of body and limbs. a. Fast green/SO staining of the GP region of WT and *ACC1-CKO* adult mice after intraperitoneal tamoxifen injection at P45 stage, b. Schematic representation of the experimental design for studying GP development in WT and *ACC1-CKO* mice at different stages, c. Representative image of the mice (WT and *ACC1-CKO*) at stage P14, P28 and P120 demonstrating the difference in their body size, Representative images showing direct measurements of tibia and femur bone at (d) P28 and (e) P120 using a digital caliper in WT and *ACC1-CKO* mice, f. Fast green/SO staining of the GP region of WT and *ACC1-CKO* to demonstrate the difference in chondrocyte organization; the yellow dotted square is zoomed to visualize the chondrocyte organization in the RZ, PZ, and HZ of the GP. Bar graph showing relative difference between (g) width of different zones and (h) the no. of chondrocytes in the GP (RZ, PZ and HZ) of WT and *ACC1-CKO* mice (n=3), i. Immunohistochemical staining with anti-COLX antibody in both WT and *ACC1-CKO* GP region, j. Bar graph showing the ratio of COLX +ve chondrocytes to total chondrocytes in three different zones (RZ, PZ and HZ) of the GP (n=3), k. Representative image obtained after the 3D µCT reconstruction of the proximal tibial trabecular region of the WT and *ACC1-CKO* mice at stage P28 and P120, l-n. Quantitative graphs comparing trabecular bone morphometry parameters between WT and *ACC1-CKO* mice at P28 and P120: (l) bone volume-to-total volume ratio (BV/TV), (m) trabecular thickness (Tb.Th), and (n)bone surface-to-bone volume ratio (BS/BV), Immunohistochemical staining with (o) anti-osterix (OSX); P28 stage and (s) anti-osteocalcin (OCN); P120 stage antibody in the proximal side of the long bone of WT and *ACC1-CKO*, p. Quantification graph showing the ratio of OSX^+ve^ cells to bone perimeter, q. Trap staining in EDTA treated paraffin section from the proximal side of the long bone of WT and *ACC1-CKO* mice at P120 days, r. Bar graph showing the Receptor Activator of Nuclear Factor kappa-B Ligand (RANKL)/ Osteoprotegerin (OPG) ratio in WT and *ACC1-CKO* tissue lysates (n=3). t. von Kossa staining of the proximal tibial bone at P120, u. Representative 3D µCT reconstructions of cortical bone from WT and *ACC1-CKO* mice at P28 and P120. Data are expressed as the mean ± SEM. Two group comparisons were evaluated using unpaired Student’s t-test with Welch’s correction. The comparison between WT and *ACC1-CKO* in different GP zones were performed using two-way ANOVA followed by Šídák’s multiple comparisons test. Statistically significant values were at p<0.05.

To better understand and mechanistically explain the impact of ACC1 disruption in chondrocytes on GP activity, we performed single-cell RNA-seq assay on tibial proximal GPs obtained from P28 WT and *ACC1-CKO* mice hind limbs (Fig. 6a). Unsupervised data analysis using the Seurat V package^32^ provided a uniform manifold approximation and projection (UMAP) consisting of 27 clusters with both non-skeletal and skeletal clusters (Fig. S7g). Skeletal clusters were identified based on the markers expressed in WT cells (Fig. S7h, i). The clusters prominently expressing *Ptprc*, *Itgam*, *Acp5*, *Csf1r*, *Spi1*, *Cd33*, *Adgre1*, *Ly6g*, *Cd3e*, *Cd79a*, *Hba-a1* or *Pecam1* were collectively annotated as non-skeletal cells. The remaining clusters were retained and re-clustered as skeletal cells. The UMAP generated from re-clustered skeletal cells (9470 cells including both WT and *ACC1-CKO*) contained 21 clusters, which consisted not only of chondrocytes but also of neighboring cells such as osteoblasts, perichondrium, and bone marrow stromal cells (Fig. 6b). These skeletal subsets were annotated based on the differential gene expression profiles in WT cells (Fig. 6c, d, Fig. S7j and Supplementary excel data 7). SOX trio gene positive clusters were annotated as chondrocytes^33,34^. *Prg4* and *Creb5* gene expression was used to distinguish AC chondrocytes (Cluster 7) from GP chondrocytes^35,36^. Cluster 1, 2, 11, and 13 showed the highest expression of *Apoe* and *Clu*, supporting their annotation as resting chondrocytes^37^. However, cluster 1 also expressed *Ccnd1*, a marker associated with G1-to-S phase progression, together with low-level *C1qtnf3* expression, positioning this cluster at the transition between resting and proliferating chondrocytes^38^. Cluster 5, and 12 had consistent expression of *Ccnd1*, *Epyc*, and *C1qtnf3*, supporting their annotation as proliferating chondrocytes^37,39–41^. Clusters 17 expressed Ihh and Col10a1 with expression of *Ccnd1*, *Epyc*, and *C1qtnf3*, suggesting progression toward hypertrophic differentiation and supporting their classification as pre-hypertrophic chondrocytes. Cluster 4 was characterized by *Acan* and *Col2a1* expression in the absence of prominent *Sox* family gene expression and was therefore annotated as a perichondria population. Interestingly, cluster 19 represented a small population with low *Sox9* and *Sox4* expression but high *Col10a1* and *Col1a1* expression, consistent with a late-stage hypertrophic chondrocyte identity^42^. Clusters 0, 3, 8, 18, and 20 were classified as contaminating populations based on enrichment of immune and hematopoietic markers. Finally, clusters 6, 9, 10, 15, and 16 showed strong expression of osteoblast-lineage markers, including *Alpl*, *Sp7*, *Bglap*, and *Col1a1*, and were therefore annotated as osteoblast-lineage cells^43,44^.

**Fig. 6.**
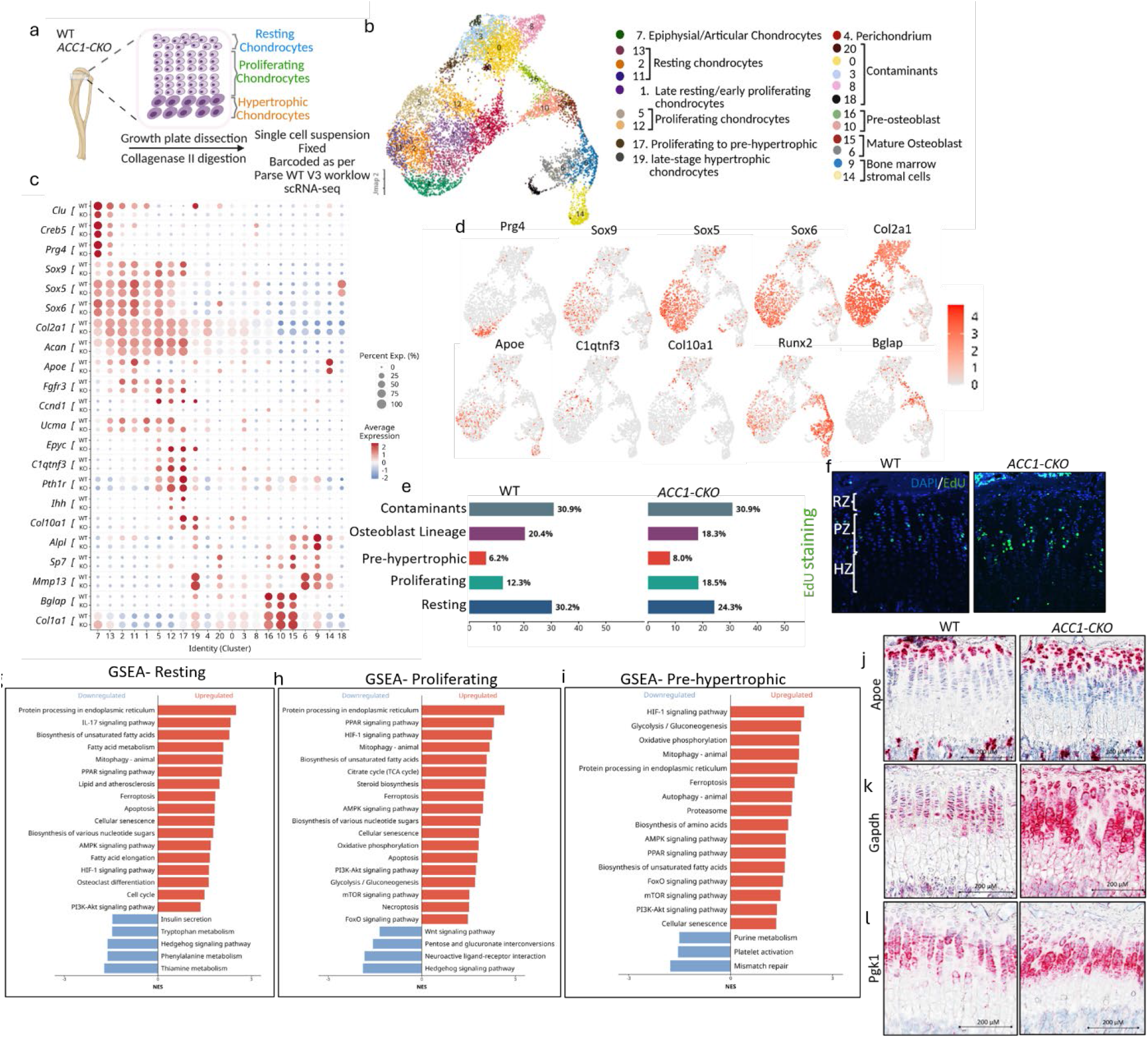
Inactivation of ACC1 in chondrocytes neonatally disrupts GP organization and activity. a. Schematic overview of the experimental setup for scRNA-seq of *ACC1-CKO* and WT GP region (for WT, n=6 and *ACC1-CKO*, n=6; each sample contained at least 5 mice), b. Integrated UMAP of 21 distinct skeletal clusters (after subsetting skeletal clusters from the original UMAP clusters containing both skeletal and non-skeletal clusters as shown in Fig. S7g) detected in scRNA-seq analysis, c. dot plot showing the expression levels of canonical markers used to identify the chondrocyte and osteoblast lineage clusters, d. UMAP feature plots color-coded to show the expression of the indicated marker genes for the skeletal subset UMAP. e. Bar graph showing the relative proportion of cells in the annotated and clubbed clusters as resting (13, 2, 11), proliferating (1, 5 and 12), pre-hypertrophic (17), f. representative image showing the increased EdU staining in the proliferating zone of the *ACC1-CKO* GP compared to WT. GSEA plot for g. resting chondrocytes, h. proliferating chondrocytes, i. pre-hypertrophic chondrocytes, In situ detection of j. Apoe, k. Gapdh, and l. Pgk1 RNAs (magenta) in tibia GP, Counterstaining, hematoxylin, 20× magnification (n=3); see supplementary datal Fig 2 for 5× magnification image.

Comparison of cellular composition revealed a prominent expansion of the proportion of proliferating chondrocytes in *ACC1-CKO* samples, increasing from 12.3% in WT to 18.5% and consistent with our histological findings, together with a modest increase in the relative amount of pre-hypertrophic chondrocytes (Fig. 6e, supplementary excel data 8). This was further confirmed by increased EdU staining in the proliferating zone of *ACC1-CKO* GP (Fig. 6f). Thus, loss of ACC1 alters the normal distribution of chondrocytes across the GP differentiation continuum. Pseudotime trajectory analysis further supported the cluster annotations, revealing a progressive transition from resting chondrocytes through late-stage hypertrophic chondrocytes and ultimately toward osteoblast-lineage cells (Fig. S8a). Analysis of genotype distribution showed an accumulation of *ACC1-CKO* cells in the middle portion of this pseudotime trajectory (inverted pseudotime 15–20), corresponding largely to proliferative-to-hypertrophic chondrocyte states (Fig. S8b, c). This shift was accompanied by stage-specific transcriptional changes, with *ACC1-CKO* proliferating chondrocytes showing upregulation of hypertrophy and terminal maturation associated genes, including *Ihh*, *Runx2*, *Col10a1*, *Mmp13*, and *Sp7*, together with altered matrix and stress-response programs (Fig. S8d).

Gene set enrichment analysis revealed distinct stage-dependent metabolic responses to ACC1 loss (supplementary excel data 9). In resting chondrocytes, ACC1 deficiency predominantly induced pathways involved in lipid handling and alternative lipid utilization, including PPAR signaling (NES-2.04, p. adjust-0.0024), fatty acid metabolism and elongation (NES-2.96, p. adjust-0.001), and biosynthesis of unsaturated fatty acids (NES-2.29, p. adjust-0.001) (Fig. 6g). This observation is confirmed by significantly upregulation of Apoe transcripts in the resting zone (Fig. 6j). These changes are consistent with a compensatory shift toward lipid scavenging and utilization when DNL is impaired. Enrichment of AMPK signaling further suggests increased energetic stress, while induction of mitophagy, ER protein processing, ferroptosis, apoptosis, and cellular senescence pathways indicates that this metabolic adaptation is accompanied by cellular stress. Notably, Hedgehog signaling was downregulated (NES--1.61, p. adjust-0.06), potentially disrupting signals required to maintain the resting chondrocyte pool and coordinate early differentiation^45^.

The response became substantially broader in proliferating chondrocytes, which face high energetic and biosynthetic demands associated with cell division and extracellular matrix production. ACC1 loss simultaneously enriched glycolysis/gluconeogenesis (NES-1.71, p. adjust-0.04), the TCA cycle (NES-2.06, p. adjust-0.009), and oxidative phosphorylation (NES-1.82, p. adjust-0.003), together with AMPK (NES-1.95, p. adjust-0.001), PI3K-Akt (NES-1.73, p. adjust-0.0001), and mTOR signaling (NES-1.52, p. adjust-0.02) (Fig. 6h). Lipid metabolic pathways, including PPAR signaling (NES-2.31, p. adjust-0.0002) and unsaturated fatty acid biosynthesis (NES-2.07, p. adjust-0.006), also remained strongly enriched. We further confirmed activation of glycolysis through RNA-Scope assays and demonstrated that transcripts for two glycolytic enzymes, Gapdh and Pgk1, robustly increased in *ACC1-CKO* GP (Fig 6k, l). This coordinated activation of multiple energy-generating, nutrient-sensing, and lipid-utilization pathways suggests a strong compensatory response to sustain the anabolic demands of the PZ in the absence of ACC1. In parallel, WNT (NES--1.63, p. adjust-0.1), and Hedgehog signaling (NES--1.81, p. adjust-0.003), were reduced, which may contribute to the disrupted differentiation and columnar organization observed in the *ACC1-CKO* GP^45^.

By the pre-hypertrophic stage, the response shifted toward a metabolic and proteostatic stress program (Fig. 6i). HIF-1A signaling (NES-2.16, p. adjust <0.0001), and glycolysis/gluconeogenesis (NES-2.07, p. adjust <0.0001) were among the most strongly enriched pathways, consistent with the increasing reliance of maturing GP chondrocytes on glycolytic metabolism. Oxidative phosphorylation (NES-2.01, p. adjust <0.0001) and nutrient-sensing pathways, including AMPK, PPAR, PI3K-AKT, and mTOR signaling, also remained enriched (NES-1.6-1.3, p. adjust<0.0001). At the same time, increased ER protein processing, autophagy, mitophagy, proteasome activity, and ferroptosis-related pathways (NES-2.01-1.8, p. adjust<0.0001) indicate increasing reliance on protein and organelle quality-control mechanisms to accommodate the metabolic and biosynthetic stress associated with ACC1 loss^46^. Together, these findings support a model in which loss of ACC1 creates a stage-dependent metabolic bottleneck: resting chondrocytes compensate through alternative lipid handling, proliferating chondrocytes broadly increase energy-producing and nutrient-sensing pathways to support their anabolic demands, and pre-hypertrophic chondrocytes increasingly engage metabolic and proteostatic stress responses^47,48^. Such sustained metabolic adaptation may ultimately compromise orderly chondrocyte maturation and contribute to the abnormal GP development observed in *ACC1-CKO* mice.

Consistent with the scRNA-seq findings, histological analyses demonstrated disrupted progression of chondrocyte differentiation across the GP zones. Type X collagen-positive cells, normally restricted to HZ, were present within the RZ and PZ of *ACC1-CKO* GPs of P28 animals (Fig. 5i, j), indicating premature or aberrant acquisition of hypertrophic characteristics. The proportion of PCNA-positive cells also increased across RZ, PZ, and HZ (Fig. S8e, f). In parallel, the numbers of TUNEL-positive cells were markedly increased in *ACC1-CKO* GPs, supporting increased cell death and complementing the enrichment of apoptosis- and ferroptosis-related pathways identified by GSEA (Fig. S8g, h). Matrix-associated changes were also evident, with significantly lower aggrecan staining in *ACC1-CKO* GPs, whereas overall SOX9 and type II collagen staining were not significantly altered, although both showed a downward trend (Fig. S8i-n). Together, these findings indicated that ACC1 loss disrupts the balance among proliferation, maturation, survival, and matrix maintenance, resulting in premature hypertrophic differentiation and impaired orderly progression of GP chondrocytes from RZ to the PZ and HZ.

We next asked whether these developmental defects extended to trabecular bone. Micro-CT analysis of the diaphysis of the long bone region showed no major genotype-dependent differences at P28. However, by P120, *ACC1-CKO* mice of both sexes exhibited reduced bone volume fraction (BV/TV) and trabecular thickness relative to WT controls (Fig. 5k, l and m). In contrast, trabecular number and separation were not significantly altered (Fig. S8o, p), indicating a trabecular thinning phenotype rather than widespread trabecular loss or perforation. Consistent with this interpretation, bone surface-to-volume ratio was increased in *ACC1-CKO* mice (Fig. 5n). This trabecular phenotype was accompanied by a reduced number of cells positive for osteoblast markers osterix (Fig. 5o, p) and reduced amount of bone-specific osteocalcin matrix protein (Fig. 5s), increased level of TRAP activity in the trabecular region (Fig. 5q), and an increased RANKL/OPG ratio (Fig. 5r), together supporting enhanced osteoclastogenic drive and reduced osteoblast differentiation. Von Kossa staining further showed reduced mineralization in *ACC1-CKO* trabecular bone (Fig. 5t). We did not observe changes in cortical bone (Fig. 5u). Thus, although ACC1 deletion does not markedly impair early postnatal trabecular bone acquisition, it compromises the long-term maintenance and mineralization of trabecular bone.

### Pharmacological ACC inhibition attenuates obesity-exacerbated PTOA

Finally, we investigated whether pharmacological targeting of ACC1 with ND-630 could serve as a therapeutic intervention to attenuate OA progression in an obesity accelerated post-traumatic OA (PTOA) animal model. Male C57BL/6J mice were maintained on either LFD or HFD to establish distinct metabolic conditions, followed by vehicle or ND-630 treatment for 28 days and non-invasive ACL rupture (Fig. 7a). As expected, HFD-fed mice exhibited substantially greater weight gain (median difference of 18.6g) before treatment than LFD-fed mice, with no differences between animals subsequently assigned to vehicle or ND-630 groups (Fig. 7b). ND-630 modestly limited to additional weight gain during the treatment period in HFD-fed mice, although the obese phenotype remained evident (Fig. 7c).

**Figure 7:**
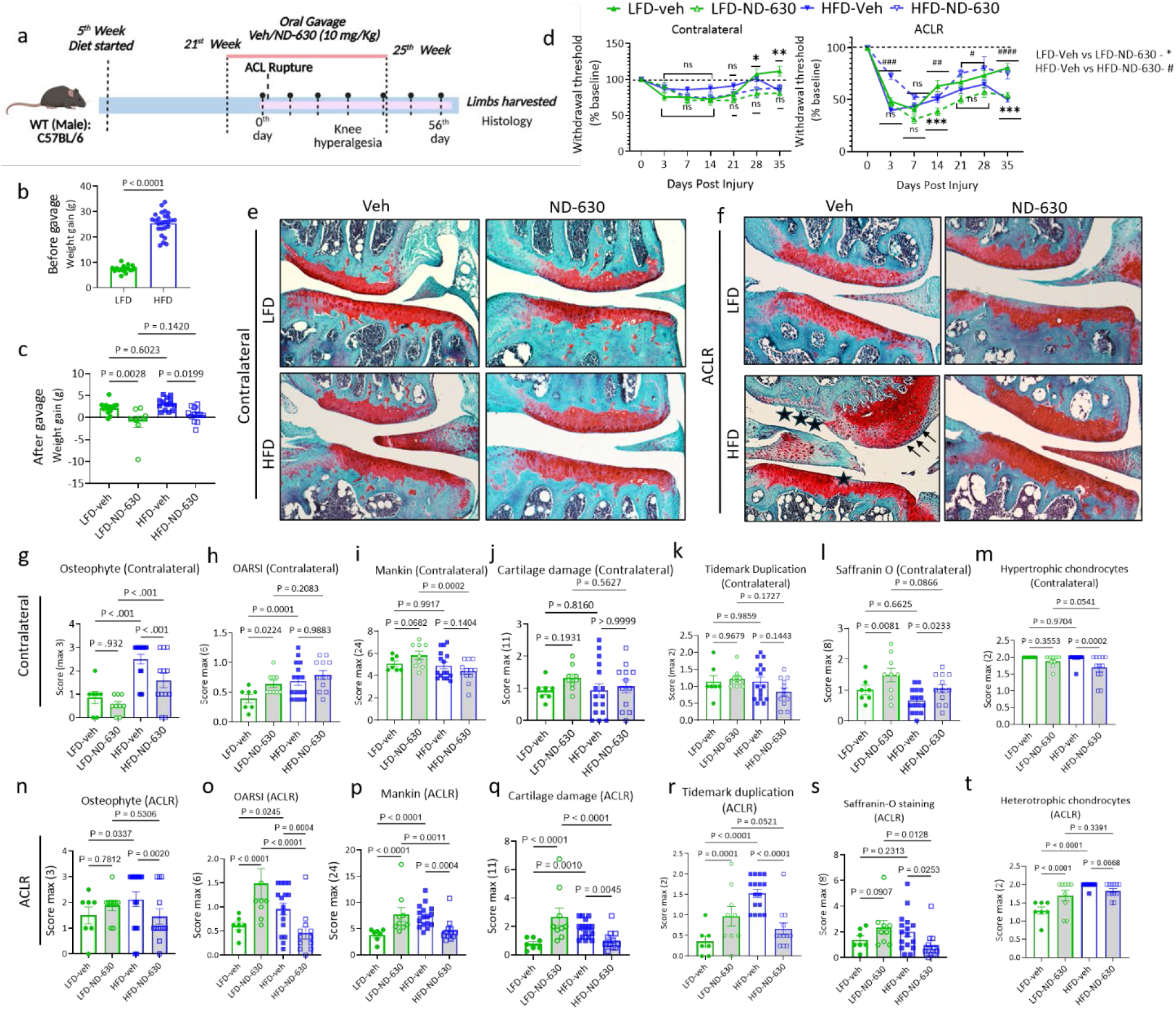
Pharmacologic ACC inhibition with ND-630 mitigates obesity accelerated post-traumatic OA. a. Schematic representation demonstrating the experimental design and timeline for obesity induction, ND-630 administration (10mg/Kg body weight) followed by non-invasive ACL rupture and assessment of post-traumatic OA development, Histogram showing the weight gained by mice in all the groups (b) before ND-630/veh gavage treatment and (c) after ND-630/veh gavage treatment, d. Line graph demonstrating the results of knee hyperalgesia analysis where withdrawal threshold (% baseline) was recorded at different time points post injury for both contralateral and ACL ruptured limb for male mice in four different groups namely LFD vehicle treated (n=7), LFD ND-630 treated (n=9), HFD vehicle treated (n=16) and HFD ND-630 treated (n=12), Fast green and Safranin O staining of contralateral (e) and ACL ruptured (f) knee joint obtained of mice from all four groups, Graphs comparing the osteophyte score, OARSI scoring, Overall mankin scoring and individual scoring parameters included in Mankin analysis for Contralateral limb (g-m) and ACL ruptured limb (n-t) among all the four groups in the study. Data are expressed as the mean ± SEM. Four-group comparisons were evaluated using two-way ANOVA followed by Tukey’s multiple comparison test. Adjusted p-values < 0.05 were considered significant.

ACL rupture induced a rapid decrease in knee withdrawal threshold in all groups, consistent with acute injury-associated hyperalgesia (Fig. 7d). In HFD-fed mice, ND-630 promoted greater recovery of withdrawal threshold during the later post-injury period compared with HFD-vehicle controls, indicating attenuation of persistent pain sensitivity. In contrast, ND-630 did not produce a comparable analgesic benefit under LFD conditions and was associated with lower withdrawal thresholds at later time points. Thus, the effect of ACC inhibition on injury-associated pain was strongly dependent on the metabolic context.

Histological examination showed relatively limited pathological changes in contralateral joints, with no consistent improvement in overall OARSI, Mankin, cartilage damage, tidemark duplication, Safranin O staining, or hypertrophic chondrocyte scores following ND-630 treatment (Fig. 7e, g–m). In ACL-ruptured joints, however, HFD markedly exacerbated cartilage degeneration, characterized by greater surface damage, proteoglycan loss, and pathological remodeling compared with LFD controls (Figure 7f). ND-630 substantially attenuated these changes in HFD-fed mice, resulting in significantly lower OARSI (mean difference of 0.61 units) and Mankin scores (mean difference of 2.82 units), less osteophyte formation (mean difference of 0.87 units), reduced cartilage structural damage (mean difference of 0.91 units), Safranin O loss (mean difference of 1.17 units), and tidemark duplication (mean difference of 0.89 units) compared with HFD-vehicle controls (Fig. 7n–s). However, hypertrophic chondrocyte scores were less consistently affected (Fig. 7t). Overall, the results indicate that ACC inhibition preferentially mitigated structural components of PTOA. Notably, the protective effect of ND-630 was most apparent in the metabolically stressed HFD condition. Under LFD conditions, ACC inhibition did not consistently improve histological outcomes and increased selected cartilage pathology measures following injury.

Together, these findings demonstrate that pharmacological ACC inhibition partially phenocopies the protective effects of genetic ACC1 suppression and preferentially attenuates obesity-amplified PTOA. These results further support ACC1-dependent metabolism as a therapeutically targetable pathway in OA while suggesting that the efficacy of ACC inhibition may depend on the underlying metabolic state.

## Discussion

Cartilage tissue architecture and metabolic homeostasis are tightly coupled, yet how chondrocytes adapt their lipid metabolic programs across differing physiological demands and disease states remains poorly understood. Our findings demonstrate that ACC1, the rate-limiting entry enzyme for DNL, functions as a stage-specific metabolic gatekeeper. While genetic deletion of ACC1 in adult AC or therapeutic inhibition of ACC *via* ND-630 drives an adaptive metabolic rewiring that confers robust protection against OA, genetic ablation of ACC1 severely compromises GP development and endochondral cartilage architecture. Recent findings from different groups align with our data that limiting excess fatty acid synthesis and lipotoxicity slows OA progression across multiple joint tissues^11,49,50^, however, we are the first to observe this striking phenotype dichotomy in GP development with ablation of DNL, suggesting a fundamental evolutionary trade-off.

We also demonstrate that by depleting cytosolic malonyl-CoA through ACC1 inhibition, global protein malonylation is reduced, thereby removing inhibitory constraints on key metabolic enzymes. This demalonylation significantly boosts the catalytic activity of enzymes across glycolysis, PPP, and the TCA cycle. This enzymatic activation triggers a robust compensatory reliance on glycolysis to meet cellular ATP demands in the absence of DNL^51^. To sustain high matrix protein synthesis and processing under lipid-depleted conditions, ACC1-deficient chondrocytes upregulate autophagic genes. We also observed a downregulation of mTOR activity (Fig. 3g, k) alongside a marked accumulation of intracellular free amino acids (Figure 2h). This coordinated upregulation of autophagic flux provides alternative energetic building blocks, allowing adult articular chondrocytes to maintain proteostasis and matrix turnover despite lipid deprivation^52^.

Despite extensive compensatory metabolic rewiring across GP chondrocytes, ACC1 deficiency ultimately leads to structural failure because this metabolic adaptation is insufficient to offset the absolute structural requirement for bulk DNL in rapidly expanding developmental cartilage^47^. At the apex of the lineage, resting-zone chondrocytes experience an energetic deficit that fails to preserve the resting stem-like pool, driven by marked downregulation of hedgehog signaling required for self-renewal and early lineage commitment. As cells transition into the proliferating zone, there is a paradoxical increase in the normalized proportion of proliferating chondrocytes supported by EdU staining and single cell analysis; however, the concurrent expression of type 10 collagen and apoptosis in these cells along with downregulation of hedgehog signaling^53–55^ indicate disruption in the crucial spatial and directional cues needed to organize these expanding cells into normal columnar stacks.

The shift toward glycolytic hyperflux and expanded TCA cycle intermediates, complemented by amino acid–driven anaplerosis^51^, increases the supply of mitochondrial reducing equivalents NADH and FADH_2_. In the context of reduced maximal respiratory capacity, this elevated electron pressure on the respiratory chain (specifically *via* succinate accumulation) would typically favor mitochondrial reactive oxygen species (ROS) generation through reverse electron transport at Complex I. However, cellular ROS levels measured *via* DCFDA remained unchanged in ACC1-deficient primary chondrocytes (Fig. S4m). We demonstrate that chondrocytes resolve this potential oxidative stress through a two-pronged redox adaptation: 1. Enhanced Antioxidant Capacity *via* the PPP: Activation of PPP enzymes significantly elevates cellular NADPH production, fueling glutathione- and thioredoxin-dependent antioxidant systems to buffer mitochondrial ROS. 2. Transcriptional Downregulation of NOX4: Transcriptomic data revealed a marked suppression of NOX4 expression in *ACC1-CKO* cartilage. By downregulating NOX4, ACC1-deficient chondrocytes in cartilage prevent the wasteful consumption of NADPH for ROS production. Instead, NADPH is conserved for antioxidant defenses, successfully insulating articular chondrocytes from oxidative damage despite extensive metabolic rewiring.

Beyond serving as the primary source of malonyl-CoA for protein acylation, ACC1 acts as the critical entry rate-limiting enzyme in DNL by converting cytosolic acetyl-CoA to malonyl-CoA. In primary chondrocyte cultures maintained under lipid-replete conditions (10% FBS), ACC1 deletion drove a depletion of total fatty acids alongside a paradoxical, unexpected elevation in saturated fatty acids (SFAs). Lipidomic profiling revealed that this metabolic shift selectively depleted major downstream fatty acid pools, including monounsaturated fatty acids (MUFAs) and polyunsaturated fatty acids (PUFAs). This persistent elevation in SFAs despite impaired DNL points toward an adaptive lipid salvage mechanism^56^. In liver-specific ACC DKO models, systemic loss of DNL triggers compensatory uptake of circulating triglycerides to maintain tissue lipid stores^57,58^. Supporting this, bulk proteomics demonstrated strong upregulation of lipid trafficking and droplet-associated proteins, including *Plin4*, *Cidec*, *Cidea*, and multiple FABP family members, indicating active lipid scavenging from surrounding microenvironments to compensate for lost endogenously synthesized lipids. Simultaneously, we observed a marked upregulation of steroid and cholesterol biosynthesis genes, suggesting that cytosolic acetyl-CoA-spared from fatty acid synthesis is actively diverted into the mevalonate pathway. This diversion aligns with the depleted free acetyl-CoA pools detected in our metabolomic analysis of ACC1-deficient cartilage. Additionally, ACC1 deletion specifically reduced ω-6 fatty acids, which are characteristically elevated during obesity and drive pro-inflammatory cascades. At the same time, ACC1 deletion enriches protective ω-3 fatty acid species in chondrocytes. We observed an elevated ratio of ether phosphatidylethanolamine to phosphatidylcholine (ether PE/PC), a distinct lipidomic signature associated with anti-inflammatory membrane dynamics, enhanced fluidity, and reduced oxidative vulnerability. In strong alignment with this anti-inflammatory lipid profile, transcriptomic analysis revealed robust suppression of inflammatory lipid signaling networks. This included marked downregulations of genes involved in arachidonic acid mobilization and eicosanoid synthesis, such as *Ptgs2* (COX-2), *Ptgs2os*, and multiple phospholipase A_2_ isoforms (*Pla2g4d*, *Pla2g4f*, *Pla2g2a*), and marked upregulation of *Pparα*, *PGC-1α*, and *Foxo6*.

In summary, our study suggests ACC1 as a stage-specific metabolic gatekeeper in cartilage biology. While indispensable during skeletal development for proper endochondral growth, ACC1 activity in adult AC promotes a lipotoxic and inflammatory environment under metabolic stress. Inhibiting ACC1 rewires chondrocyte metabolism toward a protective, demalonylation-driven glycolytic state supported by autophagic adaptation and balanced redox control. These findings highlight the critical importance of developmental and temporal context when targeting metabolic pathways, suggesting that ACC1 inhibition could be viable therapeutic strategy specifically tailored for adult OA.

Limitations: This study has several limitations. Although cartilage-specific ACC1 deletion establishes a chondrocyte-intrinsic contribution to OA, the potential roles of ACC1 in other joint tissues were not directly examined. The structural benefit of ACC1 deletion after ACL rupture was also more pronounced in females, suggesting sex-dependent responses that warrant further investigation. In addition, pharmacological efficacy was evaluated in a limited treatment paradigm and primarily in obesity-exacerbated PTOA; future studies should define optimal dosing, treatment timing, durability, and safety across broader OA contexts. Finally, although our multi-omics analyses link ACC1 deficiency to reduced protein malonylation and metabolic reprogramming, the causal contributions of individual malonylated proteins and specific lysine residues remain to be established.

## Acknowledgment

We thank the following funding support: Hevolution Foundation AGE award AGE-008 (SZ), National Institutes of Health grants R01AR081804 (SZ), R01AR086771 (SZ), R01AR082386 (SZ as a co-I), R15AR080813 (SZ), AR080062 (LV), AR83245 (LV), R01AR083966 (ZL) and R01AR086295 (ZL), Rheumatology Research Foundation Innovative Award (SZ), and Osteopathic Heritage Foundation Ralph S. Licklider, D.O. Endowed Professorship (SZ). We also thank the Small Animal Imaging Center at Yale for helping with the micro-CT analysis. We also thank Dr. Tristan Maerz and his lab (at the University of Michigan then, at ETH Zurich now) for helping us with setting up and troubleshooting the ACL-rupture platform in the lab. We thank the Chicago Center on Musculoskeletal Pain for their training in using PAM to measure knee hyperalgesia in mice. We thank Dr. Kemp E. Bruce (at the St Vincent’s Institute) and Dr. Steinberg Gregory (at McMaster University) for providing the joint samples from *ACC-DKI* mice.

## Methods

### Animals

*Aggrecan-Cre^ERT2^* mice (*B6.Cg-Acan^tm1(cre/ERT2)Crm^/J*, Stock No. 019148) and *ACC1^flox/flox^* (STOCK *Acaca^tm1Jdh^*/J, Stock no. 030954) were purchased from the Jackson Laboratory after Cryo-recovery of embryos. Cartilage-specific ACC1-deficient mice were generated by crossing *Aecan-Cre^ERT2/+^* and *Acaca^flox/flox^*mice to generate both (*ACC1^flox/flox^)* referred to as WT and homozygous mutant mice *(Acan-Cre^ERT2/+^; Acaca^flox/flox^*referred to as *ACC1-CKO).* Five-week-old C57BL/6J mice were purchased from Jackson Laboratory (Stock No. 000664) and used for ND-630 experiments. For GP study, WT dams mated with *ACC1-CKO* males received tamoxifen (100 mg/Kg of body weight prepared in peanut oil) *via* oral gavage to deliver tamoxifen to nursing pups through lactation. Both male and female offspring were genotyped. Body weight, size, and lean/fat mass were measured at P14, P28, and P120.

All the mice were housed and bred under standard conditions (22°C, 50-60% humidity, a 14h light and 10-h dark cycle) and fed a normal chow diet unless a recommended diet treatment. All the animal experimental procedures and studies were reviewed and approved by the American Association for Accreditation of Laboratory Animal Care International–accredited Institutional Animal Care and Research Advisory Committee at the Ohio University.

### Experimental design and sample size

Sample size for each experiment was provided in the figure legend. For all *in vitro* studies, a minimum of n=3 biological replicates were used. For *in vivo* experiments, sample size was determined based on a combination of several factors: availability, 3Rs for animal studies, statistical powers, and previous research experience. Based on that, we included *n* = 6 to 8 mice per group for the experiments and performed a random and blinded allocation of mice to different experimental groups.

### Chondrocyte isolation and culture

Primary mouse AC chondrocytes were isolated from either P7 or P28 WT and *ACC1-CKO* mice according to a previously published protocol^22,59^. Detailed methods are provided in the Supplemental Material.

### Diet experimental design

Both WT and *ACC1-CKO* mice (both male and female) were injected with tamoxifen (100 mg/Kg of body weight prepared in peanut oil solution) for 5 consecutive days at P45 to stimulate cre-recombinase activity to knock out the ACC1 gene in chondrocytes. A cohort of mice were fed on low fat diet (10% kcal fat; D12450Ji, Research Diets) or high fat diet or HFD (60% kcal fat; D12492i) for 25 weeks (from 8 to 28 weeks of age) to study the effect of ACC1 gene disruption on obesity induced OA in female and males. Weekly weight measurement, glucose tolerance test, body composition analysis, and knee hyperalgesia were performed on this cohort of mice (Fig. 1a). Additional detailed methods are provided in the Supplemental Material.

### PTOA experimental design

Another cohort of mice containing WT and *ACC1-CKO* (both male and female) were injected tamoxifen (IP) for 5 consecutive days at 3 months old for studying the effect on post-traumatic OA (PTOA). A non-invasive anterior cruciate ligament rupture (ACLR) was performed on the left leg of the mice using an established tibial compression-based loading system^60,61^. Briefly, the mouse (under isoflurane anesthesia) hindlimb was secured in a custom fixture that stabilized the femur while allowing axial loading of the tibia. A single, rapid axial compressive load was then applied to the tibia to generate a complete ACL rupture. Successful rupture was reliably verified through three criteria: anterior drawer laxity lacking a firm endpoint, characteristic force traces on the system’s loading-unloading graph, and an immediate post-anesthetic behavior showing avoidance of weight-bearing on the affected limb. Following spontaneous recovery, mice were returned to their home cages with unrestricted movement. The uninjured contralateral knees were used as internal controls (Fig. 1h)

### ND-630 drug interventional study in obese vs lean mice

Five-week-old C57BL/6J mice were fed a low-fat diet (10% kcal fat; D12450Ji, Research Diets) or a high-fat diet (HFD; 60% kcal fat; D12492i) till the high-fat-diet-fed animals attained more than double their initial body weight at the start of the diet treatment. Once the required body weight was attained, randomly assigned mice to vehicle and ND-630 groups in both LFD- and HFD-fed animals were administered ND-630 (10mg/Kg)/vehicle - peanut oil daily for 28 days *via* oral gavage. All the mice were kept on a special diet until the end of the experiment. The left limb of all the mice was subjected to non-invasive ACLR using the tibial compression-based loading system discussed in the previous section. The uninjured contralateral knees were used as internal controls (Fig. 7a).

### Histological analysis

Joint samples were harvested from multiple experimental cohorts: male and female WT and *ACC1-CKO* mice from the dietary intervention study; mice subjected to ACLR, (utilizing the uninjured contralateral legs as internal controls); mice from the ND-630 **d**rug interventional study; and one hind limb per mouse for GP analysis. The isolated knee joints were fixed in 4% formaldehyde for 24 hours at 4°C, followed by decalcification in formic acid (Formical-2000, StatLab) for 4 days at 4°C. Samples were subsequently dehydrated, embedded in paraffin, and coronally sectioned at a thickness of 8 µm.

Sections were deparaffinized, rehydrated, and stained with Hematoxylin, Fast Green, and Safranin-O. Histological evaluations were performed by two experienced, blinded graders (A.B. and S.M.) using randomized and coded slides. The OARSI^62^ and Mankin^63^ scoring systems were applied independently to four joint compartments: the lateral femur, lateral tibia, medial femur, and medial tibia. If scores between the two graders deviated by more than two points, the section was re-evaluated to reach a consensus. Final OARSI and Mankin scores for each joint were generated by averaging the scores across all four compartments.

### Statistical methods

All numerical results are reported as mean ± S.E.M. Statistical analysis to find the significant difference between experimental groups was analyzed by unpaired Student’s T-test with Welch’s correction, multiple t-tests without assuming equal SDs, Two-way ANOVA followed by Šídák’s multiple comparisons for pairwise comparison between WT and *ACC1-CKO* and Tukey’s multiple comparison test for four group comparisons using the GraphPad Prism software. Differences were considered statistically significant for P < 0.05. Sample size for all the experiments including in vitro and in vivo studies are mentioned in the legends.

---

**Fig. S1:**
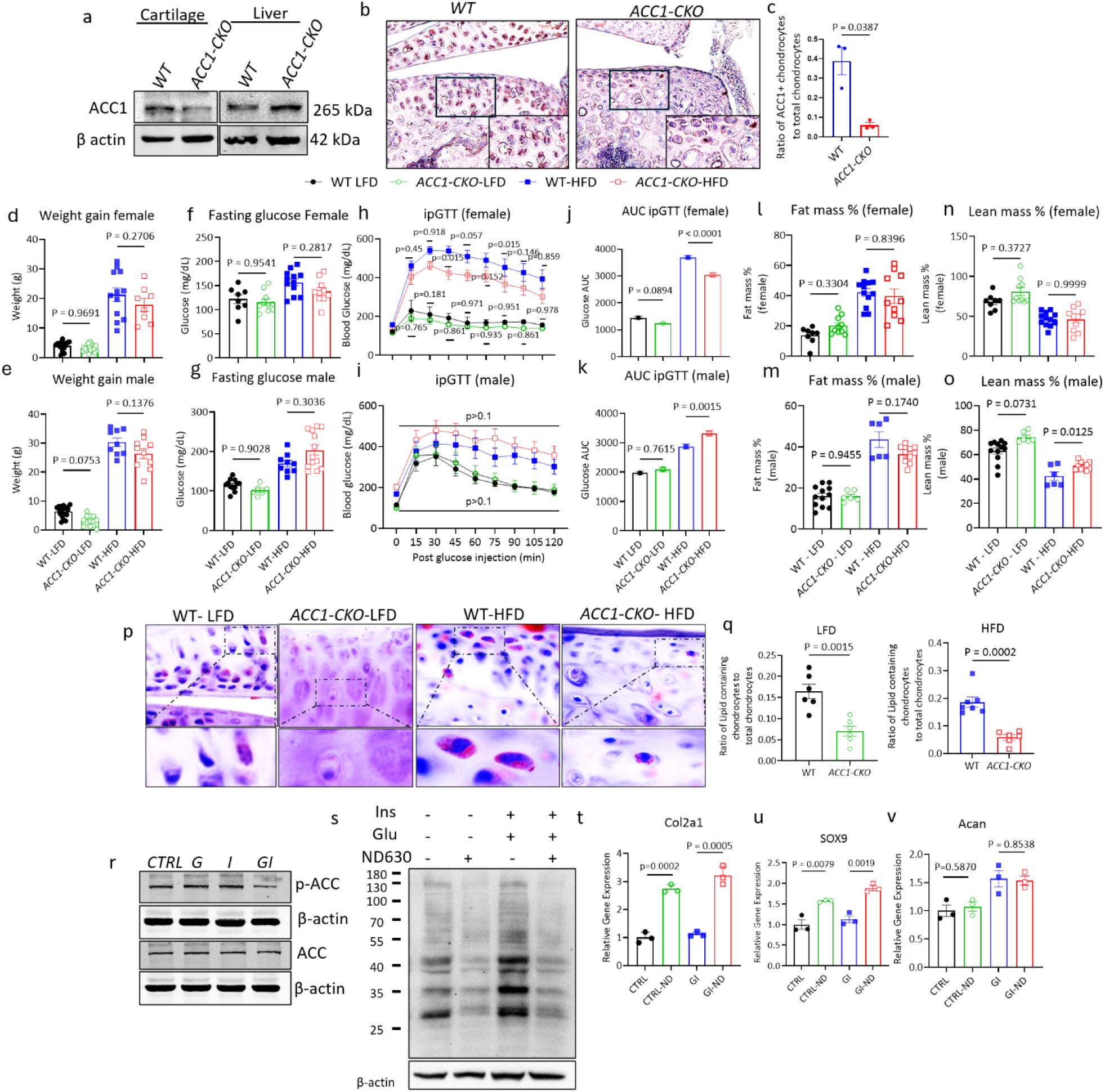
ACC activity increases in obesity conditions, and ACC1 deficiency reduces lipid accumulation in AC. a. Western blot image showing ACC1 deletion only in cartilage tissue but not in liver tissue of *ACC1-CKO* mice, n=2. b Immunohistochemical image confirming cartilage specific deletion of ACC1 in *ACC1-CKO* mice (Magnification 20×). c. Quantification of the ratio of ACC1+ chondrocytes to total chondrocytes in *ACC1-CKO* vs WT cartilage tissue (n=3). d, f, h, j, l, & n. Weight gain, blood fasting glucose (mg/dL), ip. Glucose tolerance test (ipGTT), area under the curve (AUC) for the ipGTT, fat mass %, and lean mass % in female WT and *ACC1-CKO* mice fed HFD or LFD. E, g, i, k, m, & o. Weight gain, blood fasting glucose (mg/dL), i.p. Glucose tolerance test (ipGTT), area under the curve (AUC) for the ipGTT, fat mass %, and lean mass % in male WT and *ACC1-CKO* mice fed HFD or LFD. Data are expressed as the mean ± SEM. Four-group comparisons were evaluated using two-way ANOVA followed by Tukey’s multiple comparison test. Adjusted p-values < 0.05 were considered significant. p. Oil red-O staining on the AC tissues of WT and *ACC1-CKO* mice fed on either LFD or HFD; accompanying figures below show the zoomed image of the dotted square portions marked inside each image. q. Quantification of the ratio of lipid-containing chondrocytes to total chondrocytes in WT and *ACC1-CKO* mice (n=6). r. Chondrocytes were treated with high glucose and high insulin combined or separately for 48 h (glucose (2.5/20 mM), insulin (20 nM) and were immunoblotted with anti-p-ACC and anti-ACC antibodies (n=3). s. Chondrocytes were treated with ND-630 (5 nM for 48 h) in high glucose and high insulin conditions, and the whole lysate was immunoblotted with anti-MaK antibody (n=3). t-v. Relative gene expression of cartilage ECM genes-Col2a1, Sox9, and Acan in chondrocytes treated with ND-630 in the presence and absence of high glucose and high insulin conditions (n=3). Data are expressed as the mean ± SEM. Two-group comparisons were evaluated using unpaired Student’s t-test with Welch’s correction. Four-group comparisons were evaluated using two-way ANOVA followed by Tukey’s multiple comparison test. Differences with adjusted p values <0.05 considered significant.

**Fig. S2:**
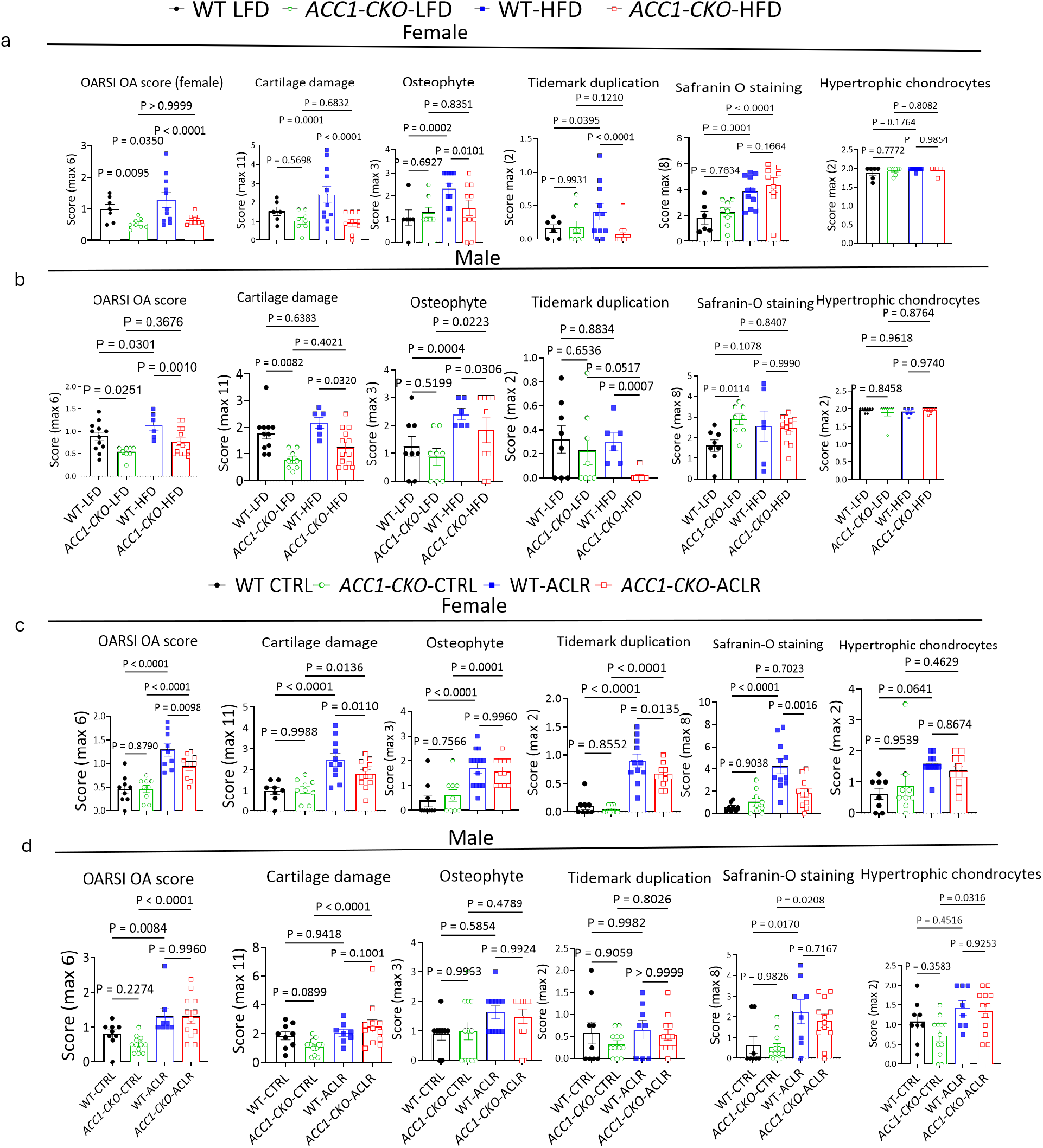
Hyperalgesia measurements and Sub-scores of the Mankin OA score for the diet and the PTOA studies. OARSI OA score and Mankin subscoring for the cartilage structural damage, osteophyte, tidemark duplication, saffranine-O staining, and hypertrophic chondrocytes observed in the fast green/Saf O stained whole knee joint obtained from female (a) and male (b) WT LFD, *ACC1-CKO*-LFD, WT-HFD and *ACC1-CKO*-HFD mice in the diet study and also female (c) and male (d) WT CTRL, *ACC1-CKO*-CTRL, WT-ACLR and *ACC1-CKO*-ACLR mice in the ACL injury study. Data are expressed as the mean ± SEM. Four-group comparisons were evaluated using two-way ANOVA followed by Tukey’s multiple comparison test. Adjusted p-values < 0.05 were considered significant.

**Fig. S3:**
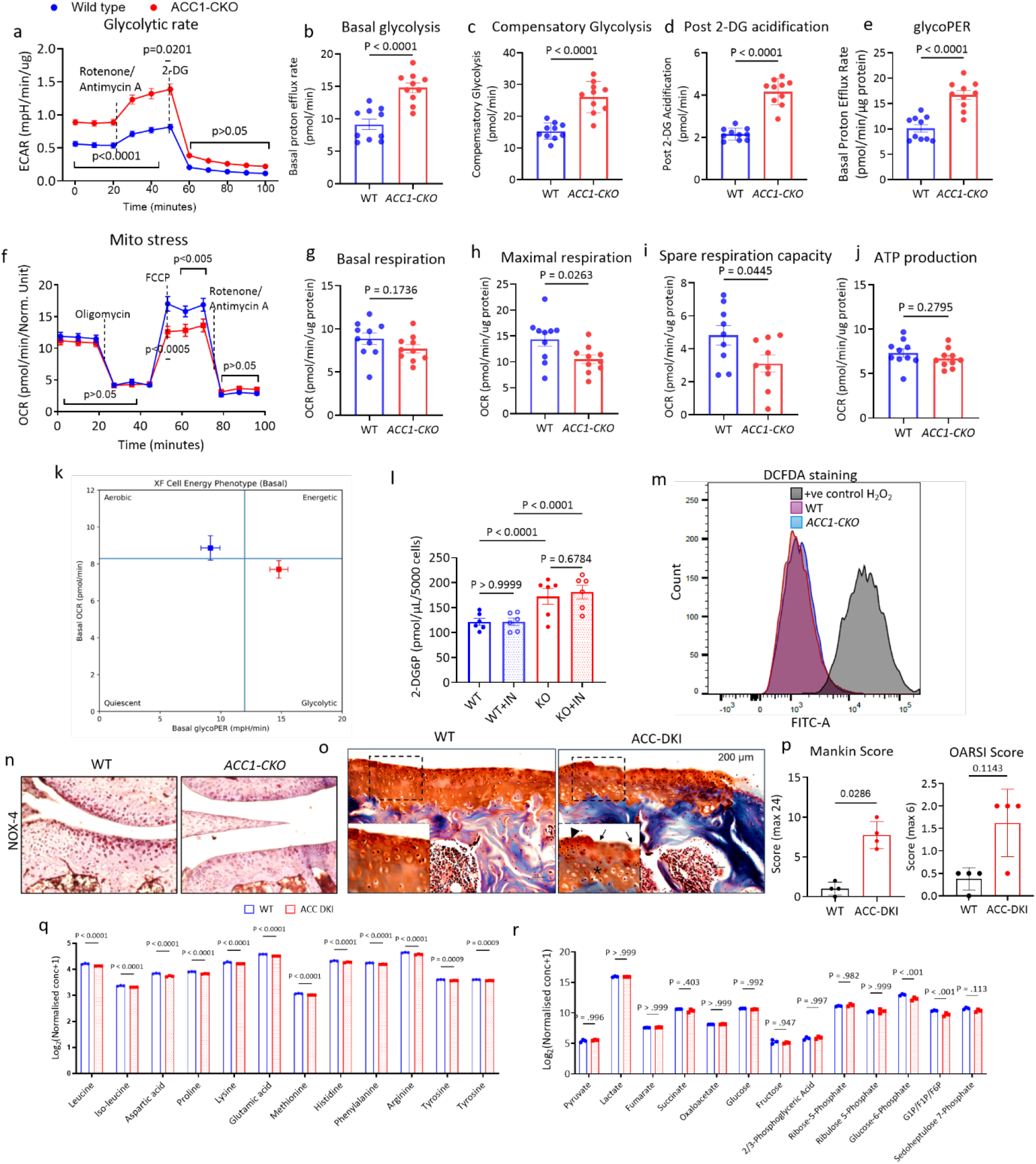
ACC1 deficiency reprograms chondrocyte metabolism towards glycolytic respiration while maintaining basal mitochondrial respiration. a. Comparison of extracellular acidification rate (ECAR) relative to function of time for WT and *ACC1-CKO* chondrocytes as measured by Seahorse Respirometry analyzer (data normalized to total protein content), (n=10), calculated respirometry parameters that compare glycolytic respiration between *ACC1-CKO* and WT chondrocytes after injection of Rotenone/Antimycin A as well as 2-deoxyglucose, b. Basal glycolysis, c. Compensatory glycolysis, d. Post-2-DG acidification, and e. glycoPER; f. comparison of oxygen consumption rate (OCR) between WT and *ACC1-CKO* chondrocytes (data normalized to total protein content), (n=10); calculated mitochondrial stress parameters between WT and *ACC1-CKO* chondrocytes; g. Basal respiration was calculated from OCR prior to oligomycin injection, h. Maximal respiration was calculated after FCCP injection, i. Spare respiratory capacity was calculated as the difference between maximal and basal respiration, j. ATP-linked respiration was determined as the decrease in OCR following oligomycin treatment, k. Seahorse energy map showing basal and stressed metabolic states of WT and *ACC1-CKO* cells. Each point represents mean OCR and ECAR ± SEM (n=10), l. 2-DG accumulation was calculated colorimetrically to measure glucose uptake in WT and *ACC1-CKO* chondrocytes with or without insulin incubation (1µM) for 20 minutes, m. Intracellular reactive oxygen species (ROS) levels were assessed by DCFDA staining followed by flow cytometry. Representative histogram (Count vs FITC-A) shows the distribution of DCFDA fluorescence intensity in untreated (WT and *ACC1-CKO*) DCFDA-stained cells and H_2_O_2_-treated DCFDA-stained cells as a positive control for ROS induction, n. Representative images of anti-NOX4 IHC in WT and *ACC1-CKO* section, o. Safranin-O staining of paraffin-embedded knee joints obtained from WT and ACC-DKI mice to analyze the status of cartilage damage, p. Mankin and OARSI scoring of the joints analyzed, n=4, q. Histogram showing fold change difference in normalized amino acid concentrations in ACC-DKI relative to WT, r. Histogram showing fold change difference in normalized glycolytic and TCA cycle intermediates in ACC-DKI relative to WT. Data are presented as mean ± SEM. Significant p-values < 0.05. Pairwise comparison between WT and *ACC1-CKO* were done using two-way ANOVA followed by Šídák’s multiple comparisons test. Two-group comparisons were evaluated using unpaired Student’s t-test with Welch’s correction. For panel q and r individual row analysis comparing WT and *ACC1-CKO* were performed using unpaired multiple t-tests without assuming equal SDs, and significant hits were determined using the two-stage step-up FDR method (*QQ* = 1%). Statistically significant value was set to p or q<0. 05.

**Fig. S4:**
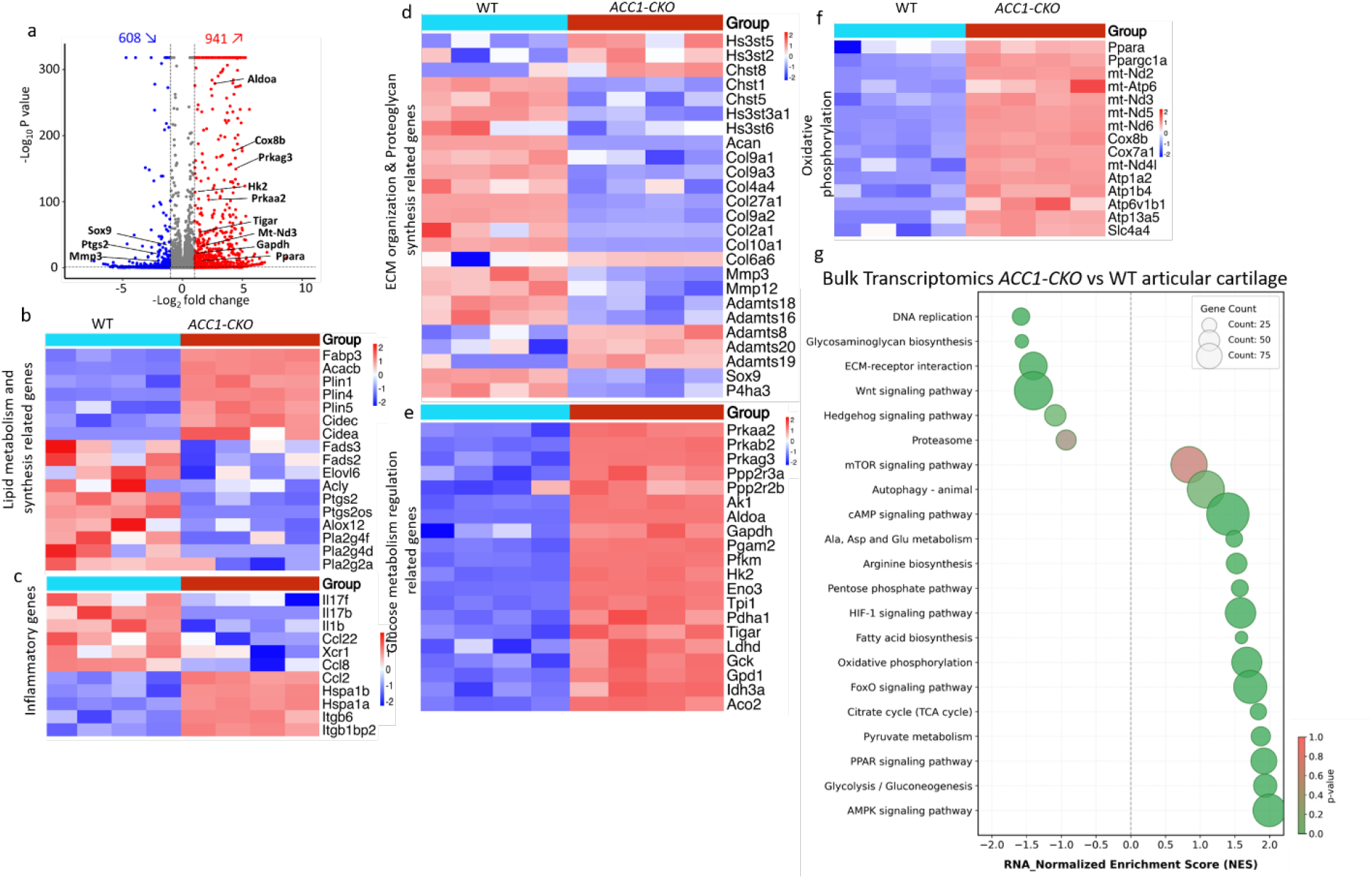
Transcriptomic analysis reveals metabolic adaptation, anti-inflammatory signaling, and extracellular matrix reorganization in ACC1-deficient cartilage. a. Volcano plot showing the differentially expressed genes in *ACC1-CKO* and WT mice AC (n=4); heatmap illustrating the differentially expressed genes between *ACC1-CKO* vs WT associated with lipid metabolism and synthesis-related genes (b), inflammatory genes (c), ECM organization and proteoglycan synthesis-related genes (d), Glucose metabolism regulation-related genes (e), oxidative phosphorylation-related genes (f), Expression values are represented as Z-score normalized by gene (red-high; blue-low). Columns denote biological replicates (n=4) and rows denote genes, clustered by expression similarity. g. Bubble plots depict significantly enriched KEGG pathways from GSE analysis of differentially expressed genes in two different groups. The x-axis shows the Normalized Enrichment Score (NES), with negative values indicating pathways downregulated in *ACC1-CKO* and positive values indicating pathways upregulated in *ACC1-CKO*. Each bubble represents a pathway; bubble size corresponds to the number of genes contributing to the enrichment, and bubble color reflects statistical significance (p-value). Pathway annotations are shown on the left.

**Fig. S5:**
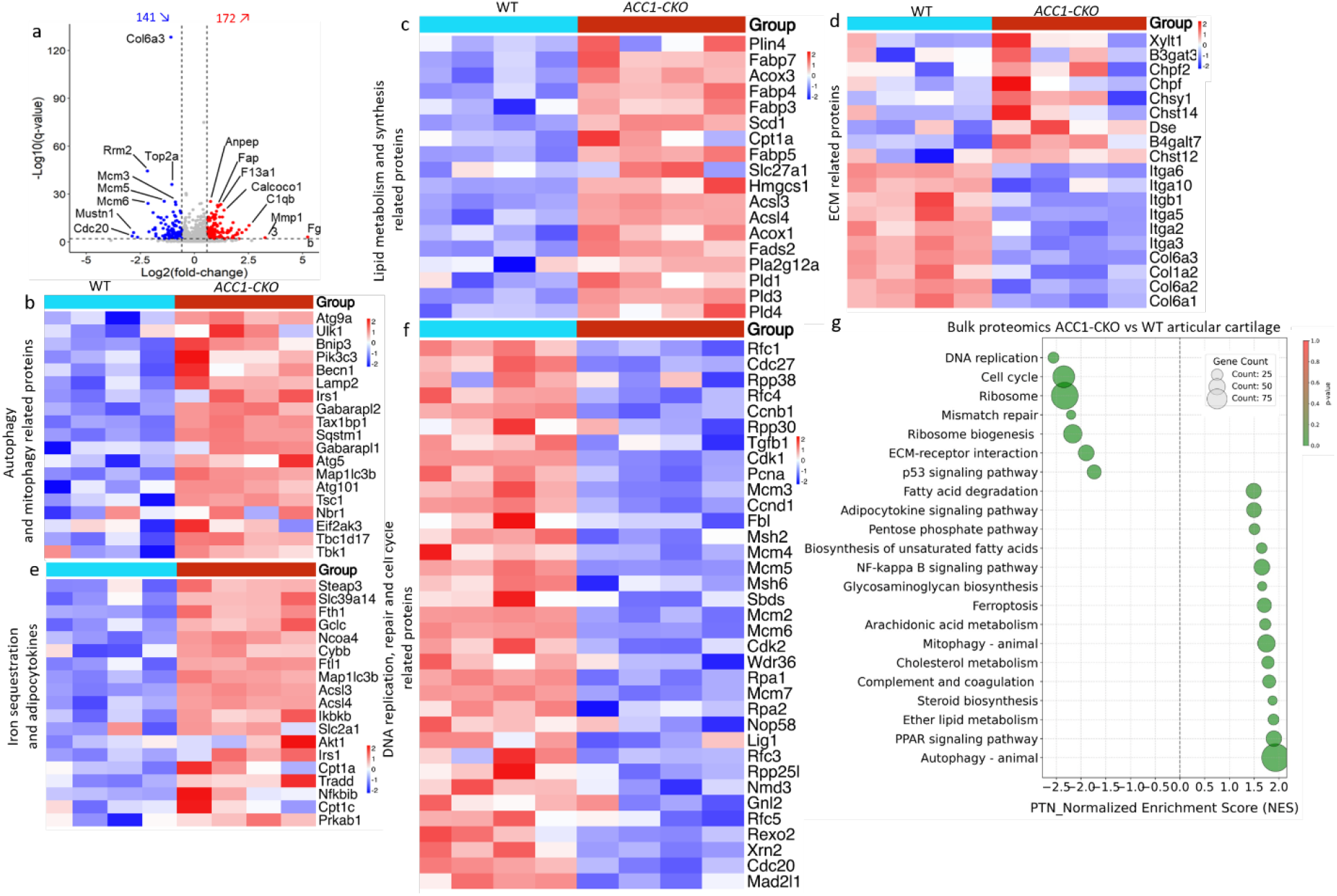
Proteomics analysis reflects an adaptive reduction in proliferative demand, by prioritizing lipid trafficking, and phospholipid remodeling. **a.** Volcano plot showing the differentially expressed proteins in *ACC1-CKO* and WT chondrocytes (n=4), Heatmap illustrating the differential expressed proteins between *ACC1-CKO* vs WT associated with autophagy and mitophagy related proteins (b), Lipid metabolism and synthesis related proteins (c), ECM organization and proteoglycan synthesis related genes (d), Iron sequestration and adipocytokines (e), DNA replication, repair and cell cycle related proteins (f), Expression values are represented as normalized Z-score with protein content in the sample (red-high; blue-low). Columns denote biological replicates (n=4) and rows denote proteins, clustered by expression similarity, g. Bubble plots depict significantly enriched KEGG pathways from GSE analysis of differentially expressed proteins in two different groups. The x-axis shows the Normalized Enrichment Score (NES), with negative values indicating pathways downregulated in *ACC1-CKO* and positive values indicating pathways upregulated in *ACC1-CKO*. Each bubble represents a pathway; bubble size corresponds to the number of proteins contributing to the enrichment, and bubble color reflects statistical significance (q-value). Pathway annotations are shown on the left.

**Fig. S6:**
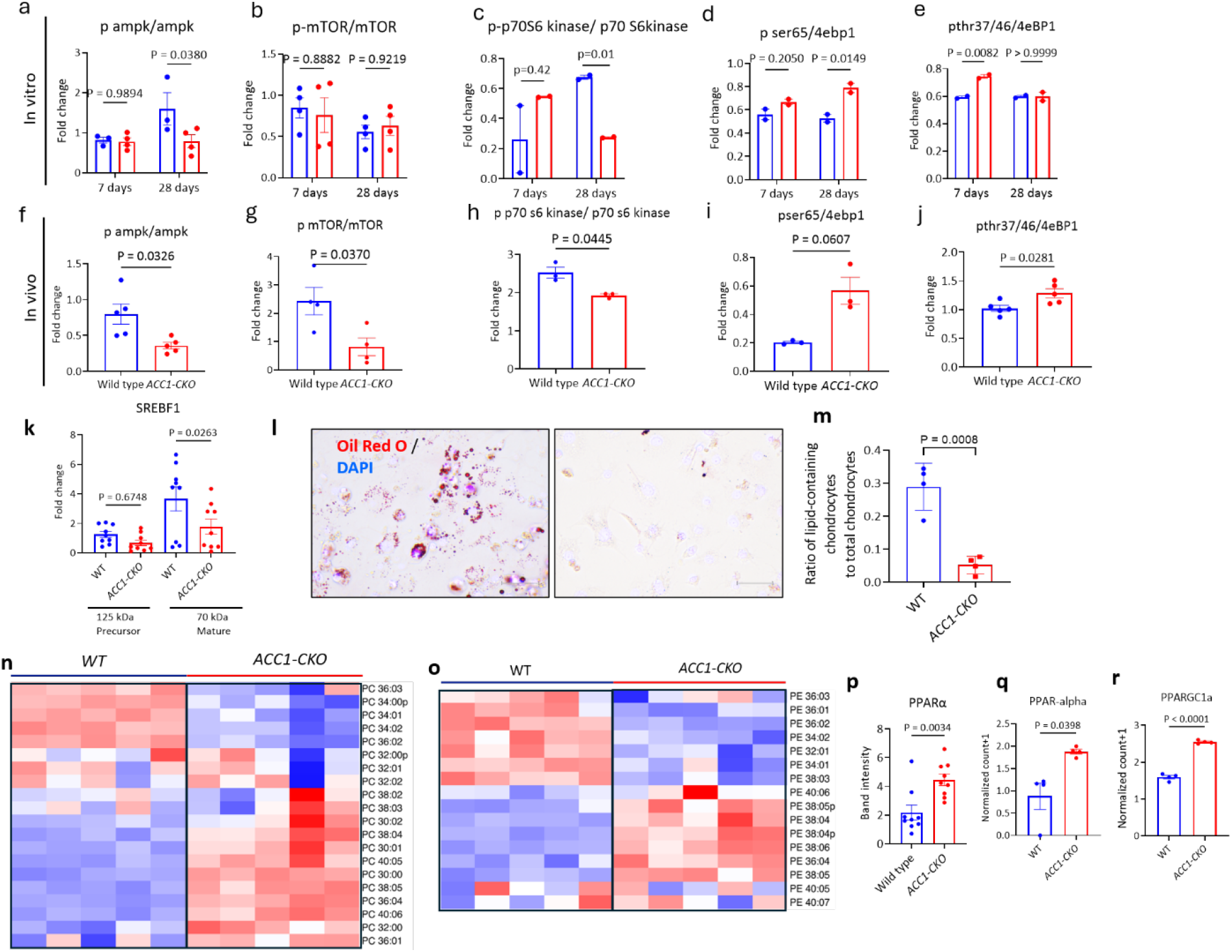
Disruption of ACC1 in chondrocytes reduces synthesis of inflammatory lipids in chondrocytes *via* PPAR-*α* signaling. Bar graph showing the fold-change difference quantified in the Western blot analysis: a. p-AMPK/total AMPK (n=4), b. p-mTOR/total mTOR, (n=4) c. p-p70 S6kinase/total p70 S6 kinase, (n=2) d. p-(ser65) 4E-BP1/total 4E-BP1 (n=2) and e. p-(thr37/46)4E-BP1/total 4E-BP1 ratios (n=2) between WT and *ACC1-CKO* chondrocytes (*in vitro*), Bar graph showing the fold change difference quantified in the western blot analysis in vivo using WT and *ACC1-CKO* AC’s homogenized lysates, f. p-AMPK/total AMPK (n=5), g. p-mTOR/total mTOR (n=4) h. p-p70 S6kinase/total p70 S6 kinase, (n=3) i. p-(ser65)4E-BP1/total 4E-BP1, (n=3) and j. p-(thr37/46)4E-BP1/total 4E-BP1 ratios, (n=5). k. Bar plot showing the relative band intensity of mature and premature SREBF1 protein in WT and *ACC1-CKO* cartilage tissue, calculated from the Western blot using ImageJ, n=9 per group. l. Oil red O staining on the WT and *ACC1-CKO* primary chondrocytes (treated with 4-HT (10µM) for 48 h), m. Quantification bar graph demonstrating the difference in lipid-containing chondrocytes (LCP) to total chondrocytes between *WT* and *ACC1-CKO* cells observed in the oil red O staining, n=4, n, o. Heat map of all detected PCs and PEs in the chondrocytes obtained from WT and *ACC1-CKO* mice p. Bar plot showing the relative band intensity of PARP-α protein in WT and *ACC1-CKO* cartilage tissue calculated from the Western blot using ImageJ, n=9 per group, q and r. Normalized count bar plot showing the expression of PARP-α and PPARGC1-α genes in WT and *ACC1-CKO* cartilage tissue, n=4. Data are expressed as the mean ± SEM. Two-group comparisons were evaluated using unpaired Student’s t-test with Welch’s correction. Pairwise comparisons were performed using two-way ANOVA followed by Šídák’s multiple comparisons test. Statistically significant value was set to p or q<0. 05.

**Fig. S7:**
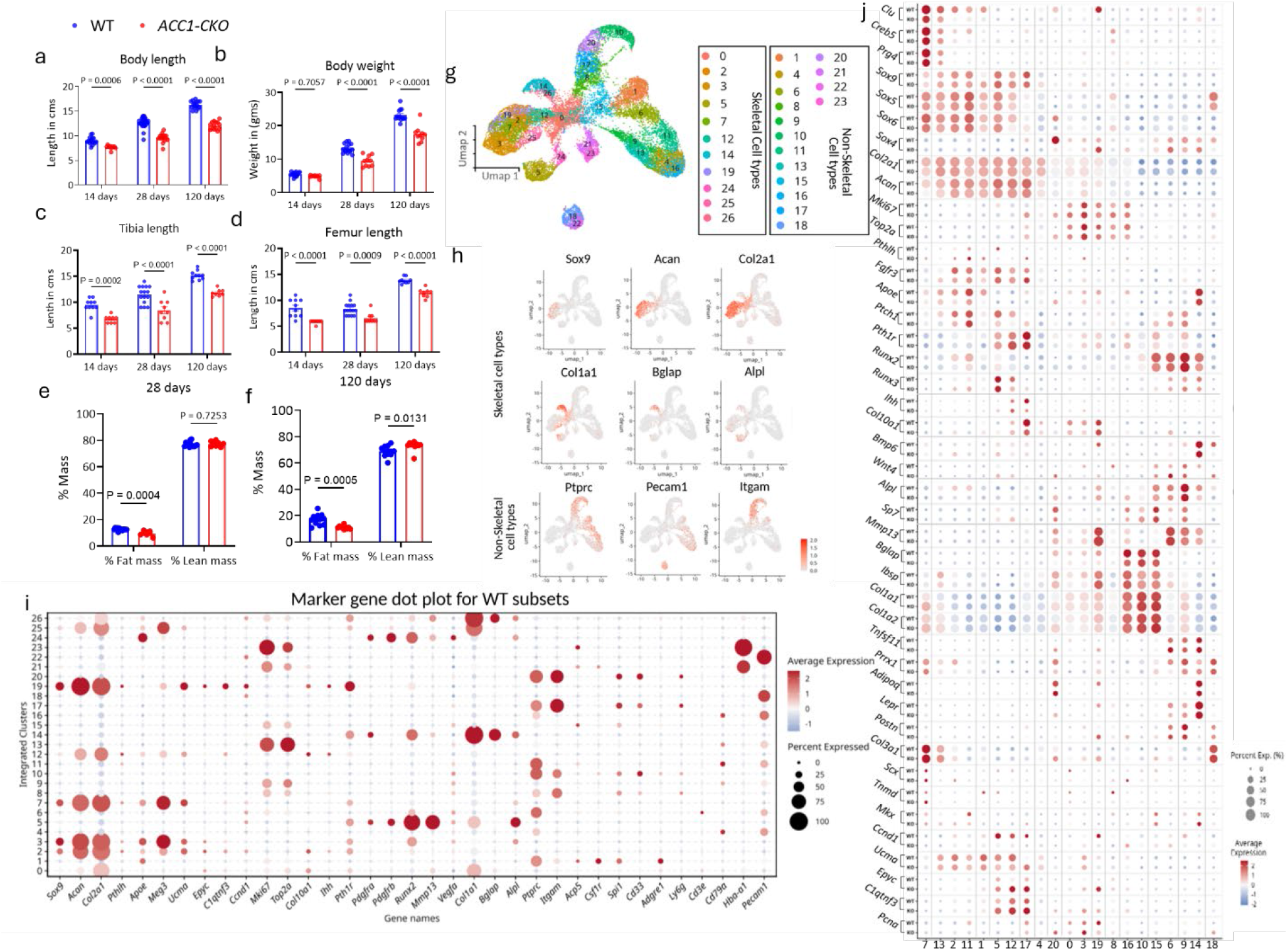
Disruption of ACC1 at early stages of development affects the growth of body and limbs. Bar graphs showing the relative change in the (a) body length, (b) body weight of WT and *ACC1-CKO* mice at different ages (for female, P14; n=16 for WT and n= *ACC1-CKO*, P28; n= 16 for WT and *ACC1-CKO* and P120; n=16 for WT and n=14 for *ACC1-CKO,* for male, P14; n=16 for WT and *ACC1-CKO*, P28; n= 16 for WT and *ACC1-CKO* and P120; n=16 for WT and n=14 for *ACC1-CKO*), Bar graph showing the relative difference between the length of (c) tibia (P14; n=8-10 for WT and *ACC1-CKO*, P28; n= 9-15 for WT and *ACC1-CKO* and P120; n=9-10 for WT and *ACC1-CKO*), and (d) femur of WT and *ACC1-CKO* mice at different ages (P14; n=8-10 for WT and *ACC1-CKO*, P28; n= 9-15 for WT and *ACC1-CKO* and P120; n=9 for WT and *ACC1-CKO*), Bar graph showing the relative difference obtained for % fat mass as well as % lean mass between WT and *ACC1-CKO* mice at stage (e) P28 (n=13 for WT and n=8 for *ACC1-CKO*),and (f) P120, (n=12 for WT and n=10 for *ACC1-CKO*), g. Integrated UMAP with 27 clusters, including both non-skeletal and skeletal clusters, from scRNA-seq analysis of digested hindlimb proximal GPs from 28-day-old *ACC1-CKO* and WT mice. h. UMAP feature plots color-coded to show the expression of the indicated marker genes. i. Dot plot showing the expression levels of canonical markers used to identify the skeletal and non-skeletal cell types (clusters were annotated based on the expression of canonical markers in WT subset), j. Dot plot showing the expression levels of extended canonical markers used to identify the chondrocytes and osteoblast lineage clusters in addition to the markers shown in Fig. 6c. Data are expressed as the mean ± SEM. Pairwise comparison were performed using two-way ANOVA followed by Šídák’s multiple comparisons test. Adjusted p-values <0.05 were considered significant.

**Fig. S8:**
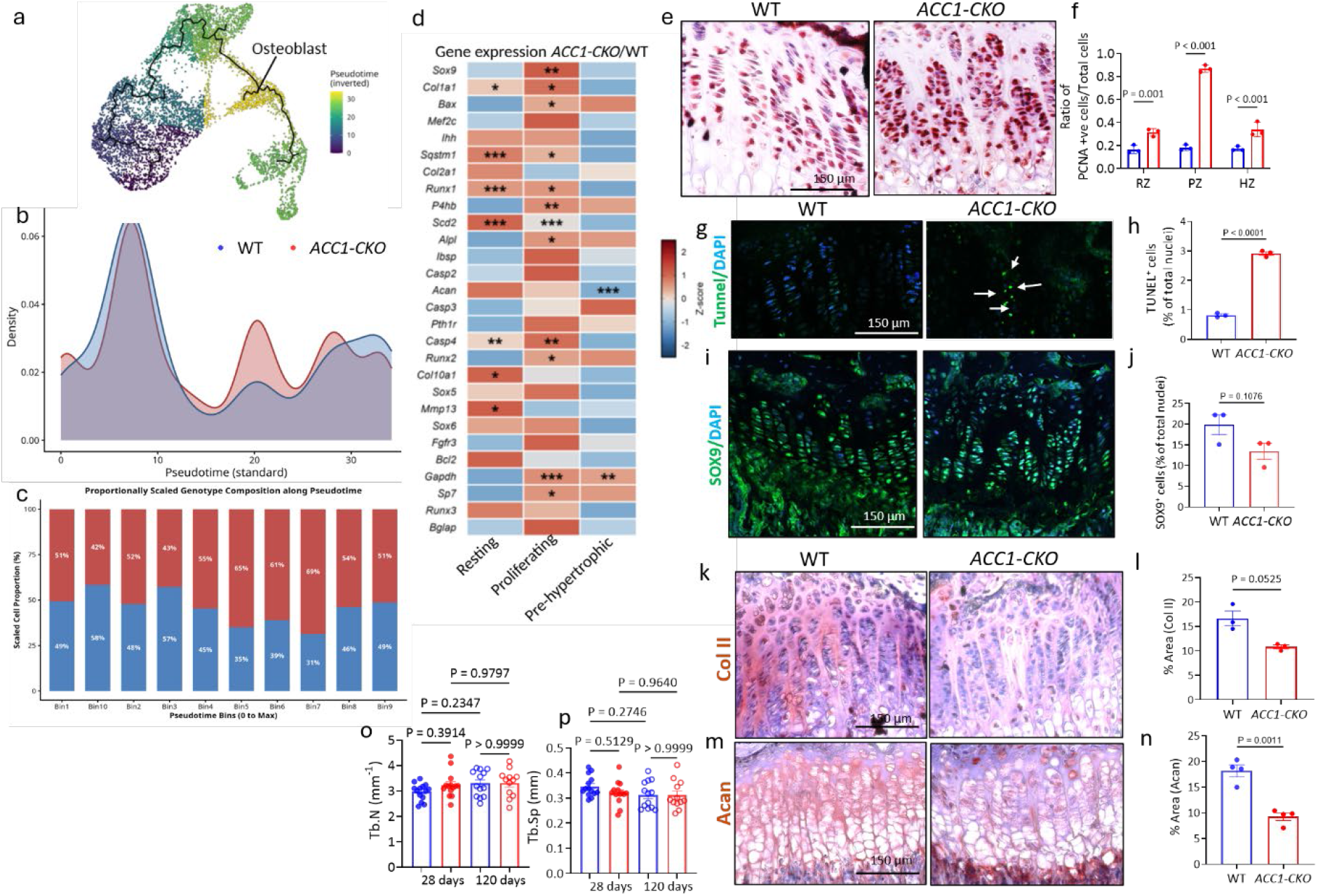
Inactivation of ACC1 in GP chondrocytes leads leads to disrupted GP development. a. UMAP displaying the inverted pseudotime trajectory, placing the developmental origin at the resting chondrocytes (purple, pseudotime = 0) and terminating at the mature osteoblast/osteocytes (yellow, pseudotime = max). This follows the traditional chronological direction of endochondral ossification. Cluster 10 was identified as the peak mature osteoblast cluster, b. Kernel density estimation (KDE) plot showing the relative distribution and frequency of cells across standard pseudotime (x-axis;0-30) comparing *ACC1-CKO* and WT cells, c. Stacked bar plot illustrating the normalized relative percentage of WT (*blue*) and *ACC1-CKO* (*red*) cells across discrete pseudotime intervals partitioned from 0 to maximum pseudotime (Bins 1–10), and d. Heatmap showing the log_2_FC in gene expression between *ACC1-CKO* and WT across defined GP zones (e.g., resting, proliferative, and pre-hypertrophic). Representative genes are categorized by functional pathways, including chondrogenesis, osteogenesis, apoptosis, and Hedgehog signaling. The color gradient indicates relative upregulation (red) or downregulation (blue) in *ACC1-CKO* relative to WT controls. e. Immunohistochemical staining with Proliferating cell nuclear antigen (PCNA) antibody in both WT and *ACC1-CKO* GP regions, o. Bar graph showing the ratio of PCNA +ve chondrocytes to total chondrocytes in three different zones (RZ, PZ, and HZ) of the GP (n=3), Representative bone sections from the proximal GP region of WT and *ACC1-CKO* mice showing (g) TUNEL immunofluorescent staining, accompanied by (h) quantitative analysis of TUNEL⁺ cells normalized to the percentage of total cells. (i) Sox9/DAPI immunofluorescent staining, with (j) quantification of SOX9⁺ cells normalized to the total cell percentage. (k) Type II collagen (Col II) immunohistochemical (IHC) staining with (l) quantification of the relative difference in Col II-positive area percentage. IHC staining for (m) Aggrecan (Acan) and (n) quantitative analysis of Acan positive area percentage. Quantitative graphs comparing trabecular bone morphometry parameters between WT and *ACC1-CKO* mice at P28 and P120: (o) trabecular number (Tb.N), and (p) trabecular separation (Tb.Sp). Data are expressed as the mean ± SEM. Two-group comparisons were evaluated using unpaired Student’s t-test with Welch’s correction. Four-group comparisons were evaluated using two-way ANOVA followed by Tukey’s multiple comparison test. Statistical significant values were set at p<0.05.

## SUPPLEMENTARY MATERIALS AND METHODS

### Chondrocyte isolation and culture

Primary mouse AC chondrocytes were isolated from either P7 or P28 WT and *ACC1-CKO* mice according to a previously published protocol^1,2^. Briefly, cartilage tissue from the femoral condyles and tibial plateau was carefully dissected from the knee joint and pre-digested in 3 mg/mL Collagenase D (Roche) in DMEM for two 45-minute periods incubation, one after the other, and then finally digested overnight at 37°C in 0.5 mg/mL Collagenase D in DMEM solution supplemented with 3 % Liberase TL (Sigma). The digested homogenate was then filtered through a 40 µm cell strainer to remove large debris and was centrifuged at 3000 rpm for 5 mins. The pellet was resuspended in complete DMEM culture medium (Life Technologies, 10567014) supplemented with 10% fetal bovine serum (FBS) and 1% penicillin/streptomycin (P/S) and cultured as passage zero cells at 37 °C and 5% CO_2_. To induce the deletion of *Acaca*, cells were treated with 4-HT (10 µM) once 70% confluency was attained. All the experiments were restricted to P1 and P2 cells.

### Glucose tolerance test

For the glucose tolerance test (GTT), mice were fasted for 8 hours and administered glucose (1.5 g/kg body weight) via intraperitoneal injection. Blood glucose levels were subsequently measured at the indicated time points using a Contour Next One glucometer (Contour).

### Body composition analysis

To analyze body fat distribution, mice were scanned using a Minispec LF50 whole-body composition analyzer (Bruker). Whole-body fat mass and lean mass were quantified and normalized to total body weight.

### Mouse knee hyperalgesia measurement

Knee joint hyperalgesia was evaluated using a Pressure Application Measurement (PAM) device (Ugo Basile, Varese, Italy), following previously established protocols^3^. Briefly, mice were restrained by the operator’s hand, and the hind paw was lightly pinned to maintain the knee in a consistent degree of flexion across all subjects. The PAM transducer was then applied to the medial aspect of the ipsilateral knee while the lateral aspect was stabilized by the operator’s thumb. Guided by the PAM software, force was applied at a constant rate of 30 g/s until a withdrawal reflex was elicited, at which point the peak force was recorded. Two independent measurements were obtained per knee and averaged to determine the final withdrawal threshold.

### Protein Extraction

Cartilage tissues from the hind limbs of WT and *ACC1-CKO* mice were dissected and homogenized in lysis buffer (150 mM KCl, 20 mM MOPS, pH 7.0) using an SI-B100 probe homogenizer. Following pulse centrifugation, the protein-containing supernatant was collected and stored at -80°C. In parallel, primary chondrocytes isolated from both groups were cultured in the presence of 4-HT and cell pellets were resuspended and incubated in IP lysis buffer, followed by centrifugation at 12,000 rpm for 10 minutes at 4°C. The resulting supernatant containing the soluble protein fraction was collected and stored at -80°C.

### Immunoblotting

Protein extracts were resolved *via* SDS-PAGE and transferred to PVDF membranes. Membranes were blocked for 30 minutes in 5% non-fat milk prepared in 1X TBST, followed by overnight incubation at 4°C with the following primary antibodies diluted in TBS blocking buffer supplemented with 0.1% Tween-20: malonyl-lysine (Cell Signaling Technology [CST], #14942, 1:500), succinyl-lysine (PTM BIO, [#PTM-401], 1:500), acetyl-lysine (CST, [#9441], 1:500), ACC (CST, [#3676], 1:1000), ACC1 (CST, [#4190], 1:1000), phospho-AMPKα (CST, [#50081], 1:1000), AMPKα (CST, [#5832], 1:1000), phospho-mTOR (Santa Cruz, [#sc-293133], 1:1000), mTOR (Santa Cruz, [#sc-517464], 1:1000), phospho-p70 S6 kinase (CST, [#9234], 1:1000), p70 S6 kinase (CST, [#9202], 1:1000), 4E-BP1 (CST, [#9644], 1:1000), phospho-4E-BP1 (Ser65) (CST, [#9451], 1:1000), and phospho-4E-BP1 (Thr37/46) (CST, [#2855], 1:1000). Membranes were subsequently incubated with IRDye 800CW or 680RD goat anti-rabbit (LI-COR, 926-68021, 1:30,000) or goat anti-mouse (LI-COR, [#926-68020], 1:30,000) secondary antibodies. Blots were imaged using the Odyssey CLx Imaging System and band intensities were quantified utilizing ImageJ software.

### Immunohistochemistry and Immunofluorescence

Deparaffinized and rehydrated sections underwent proteolytic antigen retrieval using 1× Proteinase K (from 20× stock in 1× TE) at 37 °C for 60 min in a humidified chamber. After 3 PBS washes, endogenous peroxidase was quenched with 3% hydrogen peroxide in methanol followed by blocking in 5% BSA and 0.1% Triton X-100 in PBS for 1 h at room temperature. Primary antibodies recognizing, ACC1 (Millipore sigma, [#SAB4501396], 1:100), collagen II (Millipore sigma [#MAB8887], 1:100), collagen X (Invitrogen [#14977182], 1:50) PCNA (Abcam, [#ab92552 1:100), OSX (Abcam, [#ab227820], 1:50), OCN (R&D system, (#MAB1419-SP], 1:50), Anti-NADPH oxidase 4 (NOX-4) (antibodies.com, [#AB1032], 1:100). and Aggrecan (Millipore sigma [#ab1031], 1:100) were applied and incubated overnight at 4°C. Sections were incubated for 1 h at room temperature with HRP-conjugated secondary (Goat anti-rabbit IgG horseradish peroxidase, Abcam, [ab6721], 1:500), or Goat anti-mouse IgG horseradish peroxidase, Abcam [#ab6789], 1:500). Immunoreactivity was visualized using HIGHDEF® Red HRP (ADI-950-210) counterstained with hematoxylin and mounted using a water-based medium. For SOX9 analysis, direct immunofluorescence staining was performed using a fluorophore-conjugated anti-SOX9 antibody (SANTA CRUZ sc166505, 1:50) with DAPI mounting medium confocal imaging.

### Bulk proteomics

Mouse primary chondrocytes isolated from the knee joints of P28 WT or *ACC1-CKO* mice (∼50 million cells, n = 4 each) were homogenized in a cell lysis buffer containing 0.5% SDS supplemented with 1X protease/phosphatase inhibitor cocktail (Thermo Fisher Scientific, Waltham, MA), and sonicated 2 times for 15 s at 30% amplitude using a probe sonicator. Lysates were centrifuged at 15,700 × *g* for 15 min at 4°C, after which the supernatant containing soluble proteins was collected. Proteomic data were subsequently generated from the extracts following previously established protocols ^4^. Protein concentrations were determined using a BCA protein assay (Thermo Fisher Scientific, Waltham, MA). The samples were then reduced, alkylated, and subjected to protein digestion with mini S-Trap columns. The resulting peptides were desalted using Oasis 30-mg sorbent cartridges (Waters, Milford, MA), dried under vacuum, and resuspended in 0.2% formic acid (FA) in water to a final concentration of 1 µg/µL. Prior to analysis, indexed retention time (iRT) standard peptides (Biognosys, Schlieren, Switzerland) were spiked into the samples according to the manufacturer’s instructions.^5^ LC-MS/MS analysis was performed using a Dionex UltiMate 3000 system coupled online to an Orbitrap Eclipse Tribrid mass spectrometer (Thermo Fisher Scientific, San Jose, CA). The mobile phase consisted of 2% ACN with 0.1% FA in water (Solvent A) and 98% ACN with 0.1% FA in water (Solvent B). Peptides (1 µg) were loaded onto an Acclaim PepMap 100 C18 trap column (0.1 × 20 mm, 5 µm; Thermo Fisher Scientific) at 5 µL/min for 5 min using 100% Solvent A. Separation was performed on an Acclaim PepMap 100 C18 analytical column (75 µm × 50 cm, 3 µm; Thermo Fisher Scientific) at a flow rate of 0.3 µL/min. The 210-minute gradient of Solvent B was as follows: 2% for 5 min, a linear increase to 20% over 125 min, 20% to 32% over 40 min, an increase to 80% over 1 min, a hold at 80% for 9 min, and a return to 2% over 1 min, followed by a 29-minute equilibration at 2%. Data were acquired in data-independent acquisition (DIA) mode. ^6–8^ Full MS spectra were collected at a resolution of 120,000 (AGC target: 3e6 ions; maximum injection time: 60 ms; scan range: *m/z* 350–1,650). MS2 spectra were acquired at a resolution of 30,000 (AGC target: 3e6 ions; maximum injection time: Auto; NCE: 27; fixed first mass: *m/z* 200). The DIA precursor ion isolation scheme utilized 26 variable windows covering the *m/z* 350–1,650 range with a 1 *m/z* overlap^6^. DIA data processing and statistical analysis were performed in Spectronaut (version 15.1.210713.50606; Biognosys) using directDIA. Data were searched against all *Mus musculus* protein entries extracted from UniProtKB-TrEMBL (86,521/5380 entries, 09/23/2025). Spectra were searched using Trypsin/P (up to two missed cleavages allowed), with fixed cysteine carbamidomethylation and variable methionine oxidation and N-terminal acetylation. Dynamic data extraction parameters were applied alongside non-linear precision iRT calibration and iRT profiling. Protein and precursor identifications were filtered at a 1% *q*-value, requiring at least two unique peptides per protein group. Quantification was based on XICs from 3–6 MS2 fragment ions with local normalization.

Protein abundances were log_2_-transformed and quantile-normalized, after which empirical Bayes-moderated *t*-tests were applied using the limma R package ^9^ to determine differential regulation in *ACC1-CKO* vs. WT samples. The Benjamini-Hochberg method was used to correct for multiple testing, with proteins with q-value < 0.01 and absolute log2(FC) > 0.58 defined as being differentially regulated. To identify enriched biological pathways and functions, the significantly upregulated and downregulated protein lists were used to perform Gene Set Enrichment Analysis (GSEA) using the OECloud bioinformatics platform (https://cloud.oebiotech.com). Enrichment analysis was then conducted against the KEGG pathway reference database. Pathways with a P value < 0.05 and a normalized enrichment score >1 and <-1 were considered statistically significant. Data visualization of the heat maps was also generated utilizing the OECloud platform.

### Bulk RNA sequencing

Mouse cartilage tissues were dissected from the hindlimbs of WT and *ACC1-CKO* mice (n=4 per group) and stored overnight at 4°C in RNAlater™ Solution (Invitrogen, AM7024). Tissues were homogenized using a Bead Genie bead beater in TRIzol™ Reagent (Invitrogen, 5596018). Total RNA was isolated via phenol-chloroform-isoamyl alcohol extraction followed by purification utilizing the RNA Clean & Concentrator kit (Zymo Research, R1015) and evaluated for integrity *via* gel electrophoresis. Purified samples were submitted to Novogene Corporation (Sacramento, CA) for library preparation and paired-end sequencing. Raw FASTQ sequencing reads were retrieved from Novogene and processed on the Ohio Supercomputer Center’s Owens cluster. Read quality was assessed pre- and post-trimming using FastQC (v0.11.7), with adapter and low-quality base removal performed by Trimmomatic (v0.36). Filtered reads were aligned to the GRCm39 mouse reference genome using HISAT2 (v2.1.0), followed by gene-level quantification with featureCounts from the Subread package (v1.5.0). Differential expression analysis was executed using DESeq2^10^, defining statistical significance as an absolute log₂ fold change > 0.5 and a Benjamini–Hochberg adjusted *p*-value < 0.05. To identify enriched biological functions, the significant gene lists were subjected to Gene Set Enrichment Analysis (GSEA) against the KEGG pathway reference database using the OECloud bioinformatics platform (https://cloud.oebiotech.com). Pathways were considered statistically significant if they exhibited a *p*-value < 0.05 and a normalized enrichment score (NES) > 1 or < -1. Heatmaps were similarly generated utilizing the OECloud platform.

### MaK enrichment and MS Parameter Settings

Chondrocytes from WT and *ACC1-CKO* conditions were expanded and lysed to obtain total cellular proteins, which were subsequently digested with proteases to generate peptide mixtures. The peptides were purified using C18 solid-phase extraction to remove contaminants and then lyophilized for efficient downstream processing. Malonylated peptides were selectively enriched by immunoprecipitation using anti-malonyl-lysine antibodies, enabling specific isolation of MaK-containing peptides from the complex proteome. Peptides were loaded directly onto a 50 cm x 100 µm PicoFrit capillary column packed with C18 reversed-phase resin. The column was developed with a 90-minute linear gradient of acetonitrile in 0.125% formic acid delivered at 280 nL/min.

Peptides were analyzed on an Orbitrap Fusion Lumos mass spectrometer (Thermo Fisher Scientific) utilizing electrospray ionization and HCD fragmentation. Full MS1 scans were acquired over a range of *m/z* 300–1,500 with a 3-second cycle time and a lock mass of 371.101237. MS/MS spectra were triggered with a minimum signal threshold of 500, an isolation width of 2.0 *m/z*, and a normalized collision energy (NCE) of 35.0 (Activation-Q: 0.250; Activation Time: 20.0 ms). Singly charged precursors (1+) were rejected. Dynamic exclusion was enabled with a repeat count of 1, a repeat duration of 35.0 s, an exclusion duration of 40.0 s, an exclusion list size of 500, and a mass tolerance of ±10 ppm. The resulting MS/MS spectra were processed for peptide identification and malonyl-lysine site assignment using Comet^11^ and the Harvard University Core platform. Searches were conducted against the most recent Uniprot *Mus musculus* database. Initial search tolerances were set to ±50 ppm for precursor ions and 0.02 Da for product ions. Final results were stringently filtered to a precursor mass accuracy of ±5 ppm, the presence of the intended structural motif, and a 1% false discovery rate (FDR)

### LC/MS Analysis of Metabolites

Amino acids and carboxylic acids were analyzed according to a previously reported method with modifications^12^, using an ultrahigh-performance liquid chromatography mass spectrometry (UHPLC/MS) system consisting of a Vanquish UHPLC coupled to an Orbitrap Exploris 120 mass spectrometer (Thermo Scientific, Waltham, MA, USA). Extracted samples prepared in methanol/H_2_O (8:2, *v/v*) were injected onto a BEH Amide HILIC column (2.1 × 100 mm, 2.5 µm; Waters XBridge) maintained at 40 °C. Mobile phase A consisted of H_2_O/acetonitrile (ACN) (90:10, *v/v*) containing 5 mM ammonium acetate and 0.2% acetic acid, and mobile phase B consisted of ACN/H_2_O (90:10, *v/v*) containing 5 mM ammonium acetate and 0.2% acetic acid. The injection volume was 2 µL. The separation was performed at a flow rate of 400 µL/min using the following linear gradient: 0–0.1 min, 85% B; 0.1–5 min, 85–30% B; 5–9 min, 30% B; 9–11 min, 30–85% B; and 11–20 min, 85% B. The autosampler temperature was maintained at 4 °C for all measurements. The MS detection was performed using a targeted selected ion monitoring (tSIM) method, with a resolving power of 15,000. The MS parameters were as follows: sheath gas 50 Arb units (6.35 L/min), vaporizer temperature 350 °C, spray voltage 3500 V in positive ion mode and 2500 V in negative ion mode. Amino acids were analyzed in positive ionization mode, while carboxylic acids were analyzed in negative ionization mode.

The method for the analysis of sugars and sugar phosphates was adapted from a previously published method with modifications^13^. The analysis was performed with a triple quadrupole LC/MS system (Shimadzu LCMS-8050). Separation was achieved using a Premier BEH Z-HILIC column (2.1 × 100 mm, 1.75 µm; Waters Atlantis). Mobile phase A consisted of H_2_O/ACN (90:10, v/v) containing 10 mM ammonium acetate and 0.2% acetic acid, while mobile phase B consisted of ACN/H_2_O (90:10, *v/v*) containing 10mM ammonium acetate and 0.2% acetic acid. The column temperature was set to 45 °C, and the flow rate was set to 400 µL/min with an injection volume of 20 µL. The separation was carried out using the following linear gradient: 0–0.1 min, 90% B; 0.1– 5 min, 90–85% B; 5–6 min, 85–80% B; 6–8 min, 80–45% B; 8–15 min, 45% B; 15–17 min, 85% B; and 17–20 min, 90% B. The autosampler temperature was maintained at 5 °C for all measurements. Mass spectrometric detection was performed using multiple reaction monitoring (MRM) in negative ionization mode. The MS parameters were as follows: heating gas flow 17.0 L/min, drying gas flow 3.0 L/min, desolvation line temperature 250 °C, interface temperature 400 °C, and interface voltage 4.0 kV.

### Seahorse Respirometry Analysis

Glycolytic stress assay was performed on monolayers of WT and *ACC1-CKO* P7 chondrocytes treated with 4-HT (10 µM). On the day of the experiment, the monolayer cells were incubated in XF DMEM with glucose. Basal rates of extracellular acidification were recorded over three measurement periods, then mitochondrial oxygen consumption was inhibited by injecting electron transport chain complex I and III inhibitors (Rotenone and Antimycin A, respectively), which calculates proton efflux from respiration. Further, a competitive inhibitor of glucose hexokinase, 2-DG, is injected to inhibit glycolysis, which ensures that the proton efflux rates recorded prior to the injection are primarily due to glycolysis.

Mito stress assay was performed on monolayers of WT and *ACC1-CKO* P7 chondrocytes treated with 4-HT (10 µM). These cells on the day of analysis are incubated in XF DMEM medium with pyruvate, and basal OCR was recorded over three measurement periods. Later, oligomycin injection (ATP synthase inhibitor) is introduced to calculate cellular ATP production. FCCP, an uncoupling agent was injected next to stimulate OCR and measures the maximal respiration capacity, which can also be used to calculate the spare respiratory capacity of the cells during increased energy demand or stress. The third and final injection is a combination of Rotenone A and Antimycin A, which enables determination of nonmitochondrial respiration.

### Single cell RNA sequencing

Chondrocytes were isolated from the proximal GP based on a previously described protocol with some modifications^36^. Briefly, tibia from both hindlimbs (5 mice per sample) of WT and *ACC1-CKO* mice were dissected in ice-cold complete medium (DMEM/F-12, HEPES, no phenol red, with 5% FBS and 1% Penicillin /streptomycin). AC was carefully removed by scraping using a #15 surgical scalpel and then GP was carefully dissected by making an incision between the proximal epiphysial plate and the secondary ossification center. Bone marrow cells and obvious bone tissue were removed using a blade, and the GP was stored in ice-cold 1× PBS +0.04% BSA solution in a LoBind 1.5 mL microcentrifuge tube. The tissue was then incubated and rinsed in blood cell lysis buffer (Thermo Fisher, 00-4333-57) and washed in PBS/BSA solution followed by incubation in collagenase type 2 solution. Single-cell suspension was generated by filtering through a 40-µm cell strainer into a new LoBind tube. Cell concentrations of 50,000 to 60000 cells per mL with cell viability greater than 90% were considered suitable for downstream analysis. Due to low cell yields, samples were fixed and permeabilized using Evercode Low Input Fixation v3 kits, avoiding the need for specialized equipment. Pooled cells with >70–75% viability (enriched via magnetic beads) were resuspended in storage buffer. Libraries were generated using Parse

Bioscience Evercode WT v3 split-pool barcoding, targeting 12,500 cells per library, and sequenced on an Illumina NovaSeq X Plus. Data were processed for demultiplexing, barcode assignment, and UMI quantification. Concatenated FASTQ files were processed using the Parse Biosciences split-pipe pipeline against a GRCm39 reference genome to decode combinatorial barcodes, align reads, and generate unfiltered gene expression matrices and cell metadata.

Using Seurat v5, cells passing quality control (< 10 % mitochondrial reads, 200–5,000 detected genes (nFeature_RNA), and 500–6,000 unique molecular identifiers (nCount_RNA)) were normalized *via* SCTransform, integrated and analyzed *via* Principal component analysis (PCA). Uniform manifold approximation projection (UMAP) was generated using the first 10 principal components (for shared nearest neighbor graph construction) with clustering at a resolution of 1.5, which contained both skeletal and non-skeletal clusters. The skeletal clusters were identified using the marker genes in the WT subsets, and then the identified clusters were re-clustered with the first 8 principal components at a resolution of 1.5. The final skeletal clusters were subjected to trajectory modeling and pseudotime analysis using monocle 3 where the clusters were ordered using Osteoblast/osteocytes as an initial root node. Inter-group differential expression between *ACC1-CKO* and WT within each cluster, was assessed using pseudo bulk deseq2 analysis followed by GSEA analysis. Enrichment significance was assessed using adjusted *p value* (Benjamini-Hochberg false discovery rate correction).

### RNAscope In Situ Hybridization

RNA in situ hybridization was performed on formalin-fixed, paraffin-embedded (FFPE) knee joint sections (7 µm) from P28 mice using the RNAscope 2.5 HD Reagent Kit–RED (Advanced Cell Diagnostics, Cat. No. 322360), under RNase-free conditions with RNase-free water used throughout. Following deparaffinization and hydrogen peroxide treatment, antigen retrieval was performed. Sections were hybridized with probes against mouse Gapdh, Pgk1, and Apoe, followed by sequential signal amplification (AMP1–6) per the manufacturer’s protocol. Signal was detected using Fast Red chromogenic substrate, and sections were counterstained with Gill’s hematoxylin, blued in ammonia water, and mounted with VectaMount Aqueous Mounting Medium. Stained sections were imaged using a scanner and visualized by Aperi imagescope software (v12.4.6).

### TUNEL Assay

Apoptotic cell death was assessed using the TUNEL Assay Kit (Fluorescence, 488 nm) (Cell Signaling Technology, #25879) following the manufacturer’s guidelines. Paraffin-embedded sections were deparaffinized in xylene and rehydrated through graded ethanol solutions to water. Antigen retrieval was then performed in a humidified chamber using 1X proteinase K at room temperature for 1 h, followed by a brief equilibration step as recommended by the kit protocol. The TUNEL reaction mixture was freshly prepared immediately before use by combining 1 µL TdT enzyme with 50 µL TUNEL reaction buffer per section. Samples were incubated with the reaction mixture in a humidified, light-protected chamber at 37 °C for 60 min. Sections were subsequently washed three times in PBS-TB (1× PBS containing Triton X-100 and BSA, pH 7.4). Nuclei were counterstained and sections were mounted using an antifade fluoromount-G with DAPI (Invitrogen # E142380). Fluorescent images were acquired using confocal microscopy.

### Von kossa staining

Von Kossa staining was performed on undecalcified frozen sections from 1-month-old mice to assess mineralized bone. Following dissection, samples were embedded in OCT compound without decalcification, briefly equilibrated at −80 °C for 15 min, and stored at −20 °C until sectioning. Tissues were cryosectioned at 12 µm thickness, and sections were collected using the CryoJane Tape-Transfer System onto adhesive-coated slides to preserve bone architecture. Sections were rinsed in Milli-Q water and incubated with freshly prepared 1% silver nitrate solution under a 100-W light source for 10 min. After washing in deionized water, unreacted silver was removed by incubation with 5% sodium thiosulfate for 5 min. Nuclei were counterstained with Nuclear Fast Red (Electron Microscopy Sciences #2621203) for 5 min at room temperature. Sections were then rinsed with Milli-Q water, dehydrated, mounted with a compatible mounting medium, and coverslipped for imaging.

### Trap staining

Following fixation in 4% paraformaldehyde, tissues were decalcified in 10% EDTA (pH: 7.4) for 4 weeks, dehydrated through graded ethanol, and embedded in paraffin. Sections were incubated with TRAP substrate solution (freshly prepared by dissolving 0.01% (w/v) in naphthol AS-MX phosphate substrate mix) in a humidified chamber at 37 °C for 1 h, then rinsed thoroughly with DI water before mounting.

### ELISA for determining the serum concentration of RANKL and OPG

Immediately after euthanasia, whole blood was collected by cardiac puncture using a 1 mL syringe with a 25-gauge needle and transferred into 1.5 mL Eppendorf tubes. Samples were kept at room temperature for 1 hour to allow clotting, then centrifuged at 3,000 rpm for 25 min at 4 °C. The serum layer was carefully collected into fresh tubes, snap-frozen and stored at −80 °C until analysis. Serum concentrations of RANKL and OPG were measured using enzyme-linked immunosorbent assay (ELISA) kits from R&D Systems (MTR00 Mouse TRANCE/RANKL/TNFSF11 Quantikine ELISA Kit), (MOP00 Mouse Osteoprotegerin/TNFRSF11B Quantikine ELISA Kit) following the manufacturer’s protocols.

### Enzymatic activity measurements

#### GAPDH activity assay

The activity of Glyceraldehyde 3-phosphate dehydrogenase (GAPDH) was quantified using a GAPDH activity assay kit (Abcam, ab204732). P7 primary immature chondrocytes from WT and *ACC1-CKO* mice (a litter size; each group containing at least n>5 mice) were cultured in DMEM medium supplemented with 10% fetal bovine serum and 1% penicillin/streptomycin solution in presence of 10 µM 4-HT for 48 h. The cells were then resuspended in assay buffer at a concentration of 1 x 10^6^ cells per 100 µL for each group. The cells were homogenized by passing the suspension through a 25G syringe (1 mL) 10-15 times and incubated on ice for 15 minutes, followed by centrifugation at 10,000 g for 5 minutes at 4 °C. 25 µL of the supernatant was used to measure enzyme activity according to the manufacturer’s protocol. Absorbance was measured at OD 450 nm in kinetic mode using a BioTek microplate reader. A total of 10 readings were recorded over 10 minutes for two technical replicates of each sample. GAPDH activity was calculated based on the manufacturer’s guidelines.

### Aldolase activity assay

The activity of Aldolase A was quantified using an Aldolase activity assay kit (Abcam, ab196994). P7 primary immature chondrocytes from WT and *ACC1-CKO* mice were cultured and treated the same way as performed in the previous section. Briefly, after obtaining the cell pellet, the cells were washed with 1X PBS and resuspended at a concentration of 1 x 10^6^ cells per 100 µL for each group in the assay buffer provided in the manufacturer’s assay kit. Cells were homogenized by triturating the cell suspension using a 25G syringe (1 mL) 10-15 times and then incubated on ice for 10 minutes, followed by centrifugation at 10,000 g for 5 minutes at 4 °C. 25 µL of the supernatant was used to measure enzyme activity according to the manufacturer’s protocol. Absorbance was measured at OD 450 nm in kinetic mode using a BioTek microplate reader, every 2 mins for 30 mins at 37 °C. Aldolase activity was calculated based on the manufacturer’s guidelines.

### 6-phosphogluconate dehydrogenase (6-PGD) activity assay

The activity of 6-PGD was quantified using a 6-PGD activity assay kit (Abcam, ab241016). Cells from each group were resuspended at a concentration of 1 x 10^6^ cells per 100 µL in the assay buffer provided in the manufacturer’s assay kit. Cells were similarly triturated using a 25G syringe (1 mL) 10-15 times and incubated on ice for 10 minutes, followed by centrifugation at 10,000 g for 20 minutes at 4°C. 10-25 µL of the cell supernatant was used to measure enzyme activity according to the manufacturer’s protocol. Optical density absorbance was measured at OD 460 nm in kinetic mode using a BioTek microplate reader, every 2 mins for 30 mins at 37 °C. 6-PGD activity was calculated based on the manufacturer’s guidelines. One unit of 6-PGD is the amount of enzyme that generates 1.0 µmole of NADPH per min at 8.0 at 37 °C.

### ATP assay

Cellular ATP content was measured using the CellTiter-Glo® 2.0 Assay (Promega, Cat. # G9242). An ATP standard curve was generated using Adenosine-5′-triphosphate disodium salt hydrate (Thermo Scientific Chemicals, Cat# AAL1452203), with concentrations ranging from 1 µM to 0.064 nM prepared via a series of twofold serial dilutions. CellTiter-Glo® 2.0 Reagent was added at a volume equal to that of the ATP standards or samples (100µL), mixed on an orbital shaker for 2 minutes, and incubated for 10 minutes at room temperature prior to luminescence measurement.

### NADH assay

P7 primary immature chondrocytes from WT and *ACC1-CKO* mice were cultured in the presence of 4-HT for 48 h, and NAD and NADH concentrations were quantified according to the manufacturer’s instructions. Briefly, 10^5^ cells were washed in 1X PBS and then centrifuged for 10 minutes at 125 × g at room temperature. Further, the pelleted cells were homogenized in 100 µL of NAD Extraction Buffer for NAD determination or 100 µL of NADH Extraction Buffer for NADH determination. The homogenized cells were then incubated at 60°C for 5 min, and the extract was neutralized using the opposite extraction buffer. The samples were then centrifuged at 14,000 × g for 5 min at room temperature, and 40 µL of supernatant was used for the assay. The absorbance was recorded at 565 nm at 0 and 15 min, and the concentration was calculated based on the slope recorded in the NAD(H) standard assay.

### Glucose uptake assay

Glucose uptake was measured *in vitro* P7 primary immature chondrocytes from WT and *ACC1-CKO* mice. Cells at confluency were serum-starved in glucose-free DMEM medium for 4 h prior to treatment and then treated with 1 μM insulin for 20 min at 37 °C to stimulate glucose uptake. Post-treatment, cells were washed in Krebs-Ringer-Phosphate-Hepes (KRPH) buffer (20 mM HEPES, 5 mM KH_2_PO_4_, 1 mM MgSO_4_, 1 mM CaCl_2_, 136 mM NaCl, 4.7 mM KCl, pH 7.4) and lysed. Glucose uptake was quantified using an Abcam assay kit (ab136955) per the manufacturer’s instructions. Absorbance was recorded on a microplate reader at the 412 nm wavelength, with values calculated relative to a standard curve.

### Mito tracker staining

WT and *ACC1-CKO* cells were cultured on a cover slip in 6 well plates in the presence of 4-HT for 48 h for cre-activation. The cells were thoroughly washed with 1X PBS and then stained with 5 nM Mitotracker deep red FM staining solution (Invitrogen # M22426) for 5 mins in DMEM solution. The cells were again immediately washed with a warm 1X PBS solution and fixed using ice-cold 4% paraformaldehyde for 10 mins. The cells were again washed using ice-cold 1X PBS 5 times. The coverslips with stained cells were mounted on the glass slides using a water-based mounting medium. Th cells were imaged at 644/665 nm using Nikon A1R confocal microscopy under 60X magnification. The average mitochondrial aspect ratio was calculated using ImageJ software based on the protocol described elsewhere^14^.

### Micro-CT Acquisition and Analysis

Tibial bones were scanned ex vivo using a SCANCO μCT35 system (SCANCO Medical AG, Bassersdorf, Switzerland) following standard protocols for rodent bone analysis. Proximal tibial bone was dissected after euthanasia, stored at −80 °C, fixed in 4% paraformaldehyde before subsequently imaged at a voxel size of 5–10 µm, appropriate for evaluating murine trabecular and cortical bone architecture. Scans were acquired using manufacturer-recommended settings (55 kVp, 114 µA, 900 ms integration time), with voltage and current maintained within the 45–70 kVp operating range to ensure consistent image quality across specimens. Three-dimensional reconstructions were generated using SCANCO Evaluation software, and regions of interest (ROIs) encompassing the proximal tibial metaphysis for trabecular bone and the mid-diaphysis for cortical bone (avoiding the GP) were quantitatively analyzed for bone morphometry. Quantitative parameters including bone volume fraction (BV/TV), trabecular thickness (Tb.Th), trabecular number (Tb.N), and trabecular separation (Tb.Sp) were obtained for trabecular bone using manufacturer-recommended algorithms. After applying a fixed global threshold to segment mineralized bone from soft tissue, the following parameters for cortical bone were extracted: bone volume (BV), tissue volume (TV), bone surface (BS), periosteal radius, endosteal radius, cortical circumference, polar moment of inertia (pMOI), bone tissue density, and apparent density.

